# Brain fatty acid binding protein exhibits non-preferential and mutation-resistant binding towards fatty acids

**DOI:** 10.1101/2021.09.23.461245

**Authors:** I. Bodnariuc, S. Lenz, M. Renaud-Young, T. Shandro, H. Ishida, H. J. Vogel, J. L. MacCallum

## Abstract

Members of the fatty acid binding protein (FABP) family function as intracellular transporters of long chain fatty acids and other hydrophobic molecules to different cellular compartments. Brain fatty acid binding protein (FABP7) exhibits ligand-directed differences in cellular transport behavior. For example, when FABP7 binds to docosahexaenoic acid (DHA), the complex relocates to the nucleus and influences transcriptional activity, whereas FABP7 bound with monosaturated fatty acids remain in the cytosol. We used a variety of biophysical techniques to enhance understanding of ligand-directed transport. Specifically, we examine how FABP7 binds to fatty acids, including saturated stearic acid (SA), monounsaturated oleic acid (OA), and polyunsaturated DHA. We find that at 37°C FABP7 has near equivalent binding affinities for the fatty acids, while at lower temperatures, FABP7 exhibits a preference for the unsaturated fatty acids. Therefore, nuclear localization of the FABP7-DHA complex cannot be explained by binding preferences. Using NMR spectroscopy and molecular dynamics simulations, we observe that DHA uniquely affects the portal region of FABP7, which could enhance the complex’s nuclear localization. Mutations to purported critical binding residues (R126L and Y128F) have little effect on fatty acid binding, with molecular dynamics simulations revealing that the bound fatty acid can adopt binding poses that can accommodate the mutations.

**Significance:** This work studies FABP7 at physiological temperature and shows that nuclear localization of FABP7 cannot be initiated by tighter ligand interactions. Through biophysical experiments and simulations, we show ligand-dependent conformational changes, instead of binding affinities, are associated with certain biological outcomes. Extensive simulations reveal redundancy in available ligand binding conformations, which permits mutant-resistant binding. This suggests that these mutations do not affect ligand binding affinities, but changes in protein conformation and dynamics may result in disease associated cellular outcomes.

## Introduction

Fatty acid binding proteins (FABP) belong to the intracellular lipid-binding-protein family and function as chaperones that transport hydrophobic cargo within the cell (1–3). Depending on the bound cargo, this transport can influence a host of cellular functions, including cell growth and mobility, gene expression, energy storage, and lipid metabolism (1, 2, 4).

Here, we focus on brain fatty acid binding protein (FABP7), which is found in various tissues but is particularly important in the brain. High expression levels of FABP7 are associated with increased cell proliferation, which is critical for neuron migration during brain development and has been linked to more invasive tumors and poor prognosis in glioblastomas, melanomas, and kidney and bladder cancer (5, 6). Beyond its role in cell proliferation, genome-wide association studies have linked FABP7 to a variety of diseases, including schizophrenia (7) and autism (8).

FABPs bind to a diverse range of hydrophobic cargo, including long chain fatty acids (9), endocannabinoids (10, 11), phytocannabinoids (12), and organic dyes (13) (Fig. 1D). Binding to specific ligands can lead to nuclear localization and changes in gene expression (14). For example, nuclear localization occurs when FABP7 binds to polyunsaturated docosahexaenoic acid (DHA) but not when FABP7 binds to monounsaturated or saturated fatty acids despite their similar structures (5). This differential behavior is thought to arise due to DHA binding leading to unique structural changes to the protein (5). Unfortunately, an atomistic understanding of the precise ligand-dependent structural changes that lead to nuclear localization of FABP7 are not known.

**Figure 1.**
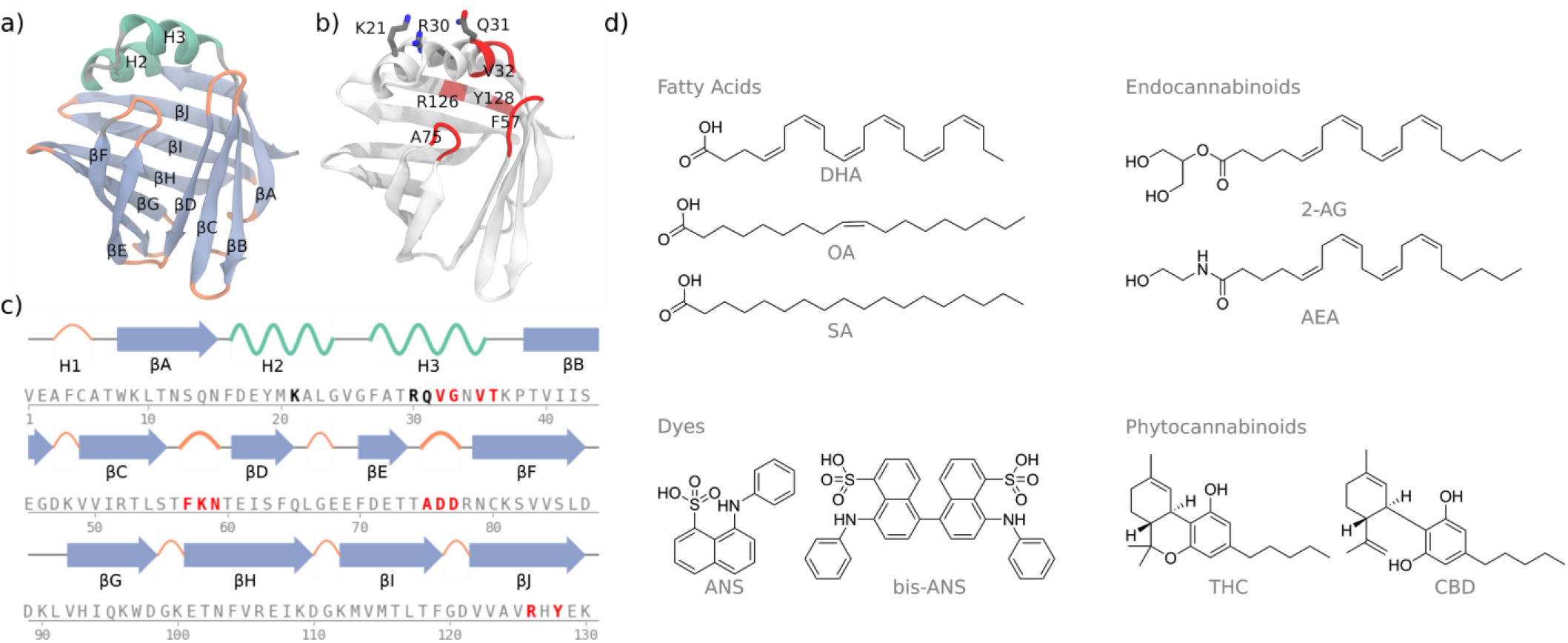
Overview of FABP7 sequence, structure, and known ligands. (a) The structure of FABP7, showing β-strands (blue), α-helices (teal), and loops (orange). (b) Key residues in the portal region including nuclear localization signal (K21, R30, and Q31). (c) The sequence and secondary structure motifs of FABP7. NLS residues shown in black and significant residues to this study in red. (d) Common ligands that bind to FABP7.

NMR and X-ray crystallographic structural studies reveal that FABPs share a common tertiary structure with a beta-barrel and flexible lid domain composed of two alpha helices (Fig. 1A-C). A common feature among FABPs is a gap between the 4th and 5th strand (denoted βD and βE in Fig. 1A). Contacts across the gap are primarily between sidechains rather than inter-strand backbone hydrogen bonds. Alongside the similar 3D structure, each FABP shares a ubiquitous and dynamic portal region (highlighted in red; Fig. 1B-C) responsible for mediating ligand entry and exit. F57 — located along the βCD turn — is responsible for opening and closing the portal region and interacts with ligands in the binding pocket (4). F57 also interacts with H3, which contains three residues (K21, R30, and Q31) that form the nuclear localization signal (NLS) (5). Binding to specific ligands may change the conformation of the NLS and promote nuclear localization. X-ray crystal structures of FABPs bound to fatty acids reveal that the fatty acids maintain similar poses in the FABP7 beta-barrel with hydrogen bonds to R126 and Y128 (15). Based on these structures, it is unclear why DHA-induced nuclear localization of FABP7 occurs and why the other fatty acids bound to FABP7 do not localize to the nucleus.

Herein, we use biophysical experiments and simulations to examine FABP7 binding to three fatty acids: oleic acid (OA), stearic acid (SA), and DHA. Our goals are to understand the interactions essential for fatty acid binding and shed light on the ligand-induced conformational changes that contribute to different biological outcomes. Our main findings are as follows:

- Both NMR and computer simulations show that the structures of the βCD, βEF, and H3 regions of apo-FABP7 are heterogeneous and dynamic, facilitating ligand entry. This heterogeneity is significantly reduced upon binding.
- Previous studies have suggested a robust selectivity for polyunsaturated fatty acids, like DHA, over other fatty acids (5, 14). In contrast, at physiological temperatures, we find only modest differences in binding affinity between polyunsaturated DHA, monounsaturated OA, and saturated SA.
- By performing experiments at multiple temperatures, we show that fatty acid binding is strongly temperature dependent. The differences between fatty acids are more considerable at lower temperatures, which is consistent with previous experimental findings (16).
- Previous literature has suggested that R126 and Y128 are key fatty acid binding residues. We find that mutations to these residues (R126L, Y128F, and R126L/Y128F) make only modest changes to ligand affinity.
- Extensive computer simulations reveal that multiple sites within the binding pocket can bind the fatty acid carboxylate group. Even wild-type FABP7 displays substantial heterogeneity in fatty acid binding poses. Upon mutation, the population of binding modes shifts to other poses that can better coordinate the carboxylate group.
- We observe that DHA binding leads to substantial chemical shift changes in nuclear localization signal residues K21, R30, and Q31. Similar changes are not observed for OA or SA.

## Methods and Materials

### Computational

#### Molecular Dynamics

The crystal structure of FABP7 bound to antinociceptive SBFI-26 (PDB ID: 5URA) (17) was used as the starting structure for our models. To simulate apo-FABP7, the inhibitor was removed. Hydrogen atoms were initially assigned with H++ (18) and adjusted to maintain hydrogen-bonding networks within the protein. The systems were solvated with a 10 Å octahedral water box and neutralized with sodium ions using the tLEaP utility from AmberTools (19).

The OpenMM (20) python library was utilized to run all molecular dynamics simulations. The Amber ff14SB forcefield (21) was used for all protein atoms. Each simulation was minimized using limited memory Broyden-Fletcher-Goldfarb-Shanno optimization, heated, and equilibrated. Production simulations were performed under NPT conditions controlled using a Monte Carlo barostat (1 bar) and Langevin thermostat (2 fs timestep and 1 ps^-1^ collision frequency). Constraints were applied to bonds involving hydrogen and the system equilibrated for 500 ns using a 2 fs timestep. For all simulations, particle mesh Ewald (PME) was applied to electrostatic interactions and all non-bonded interactions were cutoff at 10 Å.

#### Docking

AutoDock Vina (22) was used to generate docked poses of each ligand docked to structures obtained from a 1 μs simulation of apo-FABP7. Poses within 1.0 Autodock Vina scoring unit were then clustered according to the position of the binding pocket residues and the bound ligand to 30 clusters using Ward’s hierarchical agglomerative clustering algorithm, producing several unique binding poses (Fig. S1 in the Supporting Material) using the cpptraj (23) from AmberTools (19). The medioids of each cluster were used as starting structures for MD simulations. At least 50 μs of simulation time was obtained per system for a total of 800 μs of sampling time.

#### Ligand Parametrization

Each ligand was optimized at the B3LYP/6-31G(d) level of theory using Gaussian 16 (Revision C.01) (24). Following optimization, each ligand was assigned GAFF (25) atom types and AM1-BCC charges using the Antechamber utility from AmberTools (19).

#### Hidden Markov Modeling

Hidden Markov models were generated using the PyEMMA version 2.5.7 (26). The input MD coordinates were first analyzed with time-independent coordinate analysis (tICA) (27) to reduce dimensionality and isolate slowly evolving modes. These components were clustered using k-means to 100 clusters using PyEMMA default settings. The clusters were used as input to generate hidden Markov models. Implied timescale plots and Chapman-Kolmogorov tests were used to select the number of hidden macrostates and shortest lag time that generated stable models (Fig. S2). For each generated hidden Markov model, this resulted in ~2-4 states with a lag time of 10-20 ns. From the hidden Markov models, transition times and stationary distributions of each macrostate were calculated.

### Experimental

#### Cloning

FABP7 gene sequence for wt, R126L, Y128F, and R126L/Y128F gene sequences were ordered from Life Technologies in the pENTR plasmid that is compatible for GATEWAY cloning. A TEV cleavage site followed by a 4-Gly spacer was included on the N terminus each sequence. The LR clonase reaction was used to clone all sequences into pHGWA His-tagged expression vector.

#### Protein Expression, Purification, and Delipidation

Protein was expressed in *Escherichia coli* BL21 (DE3) cells and induced using 1mM IPTG. Cell pellets were resuspended in lysis buffer (25 mM imidazole, 3 mM DTT, 50 mM sodium phosphate, 250 mM NaCl, pH 7.8) and lysed with sonication. The soluble fraction was separated from cellular debris by centrifugation at 40000g. His-labelled FABP7 was then immobilized with Ni-NTA IMAC column (AKTA system, GE) and eluted in high imidazole solution (250 mM imidazole, 50 mM sodium phosphate, 250 mM NaCl, pH 7.8). The protein was incubated overnight with TEV (VWR) at 4°C and Histag and TEV was purified out with Ni-NTA beads (GE). Because FABP7 binds to hydrophobic molecules, the recombinant protein must be delipidated from endogenous bacterial lipids. This is performed by two successive incubations of FABP7 with Lipidex-5000 beads (Perkin Elmer) for one hour each at 37°C. Purified protein was then dialyzed into a final solution (20 mM sodium phosphate, 120 mM NaCl, pH 7.4). ^15^N isotopically labelled protein used for NMR experiments followed the same protocol except cells were grown in M9 minimal media with ^15^N-ammonium chloride. ^15^N-FABP7 was dialyzed into NMR buffer (20 mM sodium phosphate, pH 7.4) for all NMR experiments.

#### NMR

Spectra were collected using a Bruker 600 MHz Avance spectrometer equipped with a 1H, 15N, 13C TXI probe with a single axis z-gradient. Data processing was completed using NMR Pipe and NMR Draw and analyzed with NMR View. All spectra were collected at 25°C and referenced to sodium trimethylsilylpropanesulfonate (DSS).

^1^H-^15^N HSQC spectra was collected with a 50 µM FABP7 concentration. Fatty acids were titrated into 50 µM FABP7 sample from a DMSO stock, to a maximum of 2.5% DMSO volume content. Control spectra of FABP7 with 2.5% v/v DMSO showed no changes from DMSO free spectra. Assignment of amide backbones was taken from the literature for apo, DHA and OA bound protein complexes (28, 29). SA-bound FABP7 assignment of ^1^H-^15^N HSQC spectra was completed by peak following in the titration experiment. Ambiguous peaks from spectra were excluded from this study.

Weighted chemical shift perturbations between apo and fatty acid bound spectra were calculated as follows:

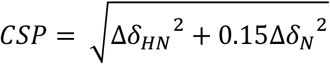

Where Δ*δ*_*HN*_ and Δ*δ*_*N*_ is the chemical shift difference between apo and holo spectra for the proton and nitrogen dimension, respectively. For residues that have multiple peaks in the apo spectra, the most intense peak was used for the CSP calculation.

#### Fatty Acids Serial Dilutions

OA, DHA, and 1-anilino-8-naphthalene sulfonate (ANS) was purchased from Sigma, and stearic acid from Larodan. Stock solutions and 1:1 serial dilution were all preformed in DMSO. Fatty acid-DMSO were mixed with assay solutions to a maximum 2% DMSO content. All dilution series were completed in low-bind Eppendorf tubes.

#### Microscale Thermophoresis

Microscale thermophoresis (MST) measurements were carried out using Monolith NT.115Pico instrument. Measurements were carried out in MST-buffer (20 mM phosphate, 120 mM NaCl, at pH 7.4, in 0.005% Tween-20) using premium capillaries (Nanotemper). Proteins were labeled with RED-tris-NHS dye following supplier protocol. Measurements were taken with instrument set to high MST power, and 20-30% LED power. A solution of 20 nM FABP7-NHS-RED-Tris labeled was combined with fatty acid dilution series. Samples were incubated for 30 minutes at measurement temperature before loading samples into MST capillaries.

Analysis of MST traces was completed using MO.Affinity Analysis v2.3 (Nanotemper) (30). Relative florescence (F_norm_) is determined using the same defined time intervals for the initial and thermo-diffusion florescence. All measurements are completed in triplicate.

#### ANS Displacement

All assays were completed at 25°C using Biotek Cytation 5 plate reader. Binding interaction measurements were performed in the same manner for FABP7 wild-type and mutant proteins. ANS signal intensity increases 40-fold when in the hydrophobic environment of the protein binding pocket. Binding affinity assay was determined using procedure previously described in literature (13).

Full experimental and computational methods are available in the supplementary material.

## Results and Discussion

### 1. FABP7 does not display a binding affinity preference for polyunsaturated fatty acids at physiological temperature

A preferential interaction has been reported between FABP7 and polyunsaturated fatty acids (e.g. DHA), over monounsaturated (e.g. OA), and saturated fatty acids (e.g. SA) (5, 16). We have performed three different binding assays (MST, fluorescence anisotropy, and ANS displacement) to characterize the affinity of DHA, OA, and SA with FABP7 at 25°C (Figs. S3-S7). We measured tighter binding affinities for DHA and OA over SA, which is consistent with the proposals that binding favors unsaturated FAs over saturated FAs (5, 16, 31).

FABP7’s preference for OA and DHA over SA largely disappears at physiological temperature (37°C). Van’t Hoff analysis from MST binding assays at 25, 29, 33, and 37°C reveals a strongly temperature-dependent binding affinity for OA and DHA, while there is little change in binding affinity for SA at different temperatures (Fig. 2). The Van’t Hoff analysis reveals that the binding of FAs is driven by enthalpy, which is consistent with previous isothermal titration calorimetry studies (16). The strong temperature-dependent binding of OA and DHA indicates that binding is entropically unfavorable.

**Figure 2.**
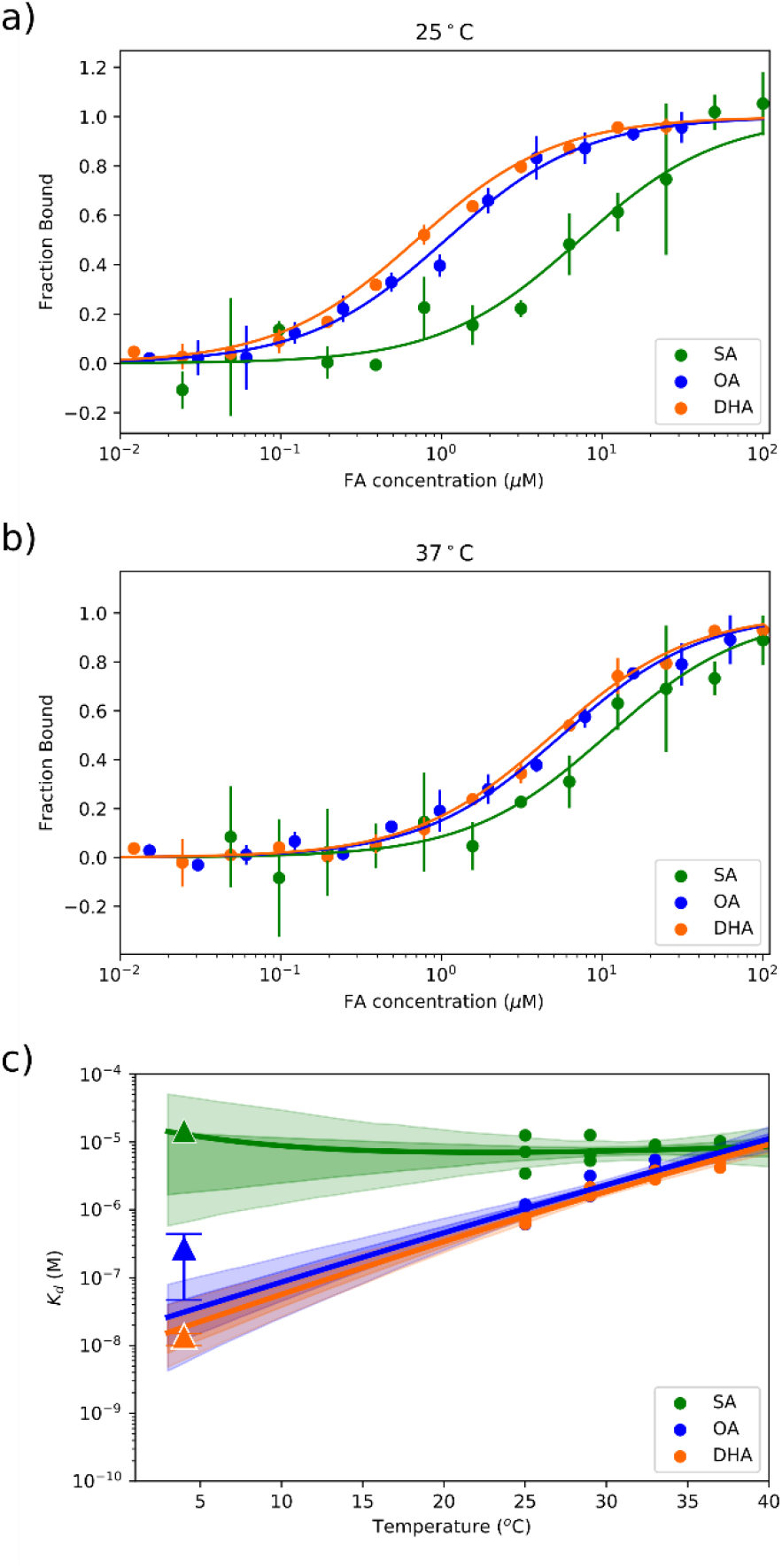
FABP7 preferentially binds OA and DHA at 25°C but not at 37°C. a-b) Binding curve measured for SA (green), OA (blue) and DHA (orange) from MST measurements at 25ºC (a) and 37 ºC (b) show preferential binding is not seen at physiological temperatures. Bars show standard deviation from triplicate measurements. c) Van’t Hoff analysis of MST binding data (circles) suggests temperature-dependent binding for OA and DHA but not SA. Literature data using the Lipidex assay (triangles) obtained at 4°C are shown for comparison with error bars indicating the spread of the literature data (16).

Previous work suggest FABP7 exhibits a strong binding preference for unsaturated fatty acids (OA, DHA) over saturated fatty acids (SA) (5, 16, 31), while our data indicates that FABP7 exhibits little selectivity at 37°C (Fig. 2). This discrepancy may arise because previous studies were based on Lipidex assays (16, 31), which include a long incubation step at 4°C. When we extrapolate our data to 4°C, we also observe a preference for unsaturated fatty acids and overall, in agreement with previous data. Our results at 37°C suggest that only modest binding affinity differences exist among fatty acids at physiological temperature.

### 2. FABP7 affinity for fatty acids is resistant to mutations of putative binding residues

X-ray crystal structures of FABP7 bound to fatty acids show that R126 and Y128 form hydrogen bonds with the bound ligand and a F104A R126A Y128A triple mutant showed no discernible binding for DHA at 37°C *in vivo* (5).

We produced and purified FABP7 with R126L, Y128F, or R126L/Y128F isosteric mutations. Our goal was to remove the ligand hydrogen-bonding ability of these residues, while minimally perturbing the structure. We expected these mutations to significantly alter the binding affinity of FABP7 towards fatty acids *in vitro*. Previous work suggests that these residues may be critical for fatty binding (15). We used ANS displacement assays to measure the binding affinity for all three mutants and MST for R126L and R126L/Y128. The thermophoretic signal was too small to determine an accurate binding affinity for Y128F using MST.

Our binding assay results indicate that the R126L, Y128F, and R126L/Y128F mutations have only modest effects on fatty acids binding. Specifically, for a given fatty acid, the binding affinity is within an order of magnitude regardless of the mutation (Figs. 3 and S4-S6). This result is surprising because the fatty acid carboxylate consistently coordinates to R126 and Y128 in X-ray crystal structures.

**Figure 3.**
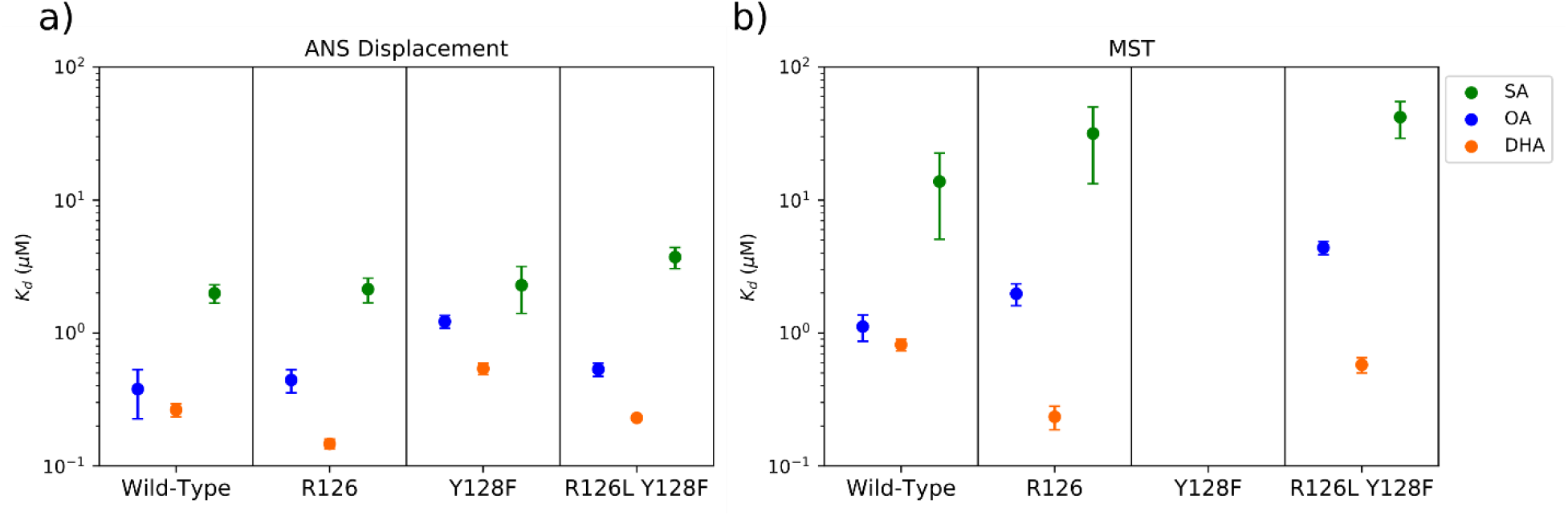
FABP7 binding affinity for fatty acid is resistant to R126L and Y128F mutations. Each panel shows the binding affinities for a particular variant of FABP7 binding to fatty acids at 25ºC: (a) ANS displacement assays, (b) microscale thermophoresis. Error bars indicate standard deviation across three replicates.

Summing our temperature series and mutagenesis binding affinity data together, it is clear that: (*i*) FABP7 does not have a strong preference between the three fatty acids tested at physiological temperature; and (*ii*) fatty acid binding is resistant to mutation of critical binding residues. The lack of specificity towards DHA is perhaps not surprising given that FABP7 can accommodate many structurally diverse ligands. These ligands, including organic dyes, endocannabinoids and phytocannabinoids, may interact with pocket residues other than R126 and Y128.

### 3. apo-FABP7 is highly dynamic with H3- and gap-unfolded conformational states

FABP7 binding to DHA leads to nuclear localization (5) and we sought to provide a structural perspective on the ligand-dependent subcellular localization of FABP7.To investigate how binding different ligands changes the structure and dynamics of FABP7, we used a combination of ligand titration protein NMR experiments and molecular dynamics simulations.

We first establish a point of reference by examining the dynamics of apo-FABP7. Our ^1^H-^15^N HSQC NMR reveal that several residues (G24, V25, G33, T36, F57, K58, N59, and T60) produce multiple peaks suggesting the presence of multiple metastable states (Figs. 4a-b and S8). Previous studies have reported that several residues within the portal region have multiple peaks for FABP7 (28, 32) and heart-FABP (FABP3) (33). Since these residues are near F57, it has been proposed that these multiple peaks observed in the apo-NMR spectra arise due to ring current effects that fluctuate with the conformation of F57. However, a F57S FABP3 mutant still produces multiple peaks for these residues (33), suggesting these peaks arise independent of ring current effects. Therefore, it is probable that different backbone conformations of portal region residues result in slow exchange and lead to this phenomenon.

**Figure 4.**
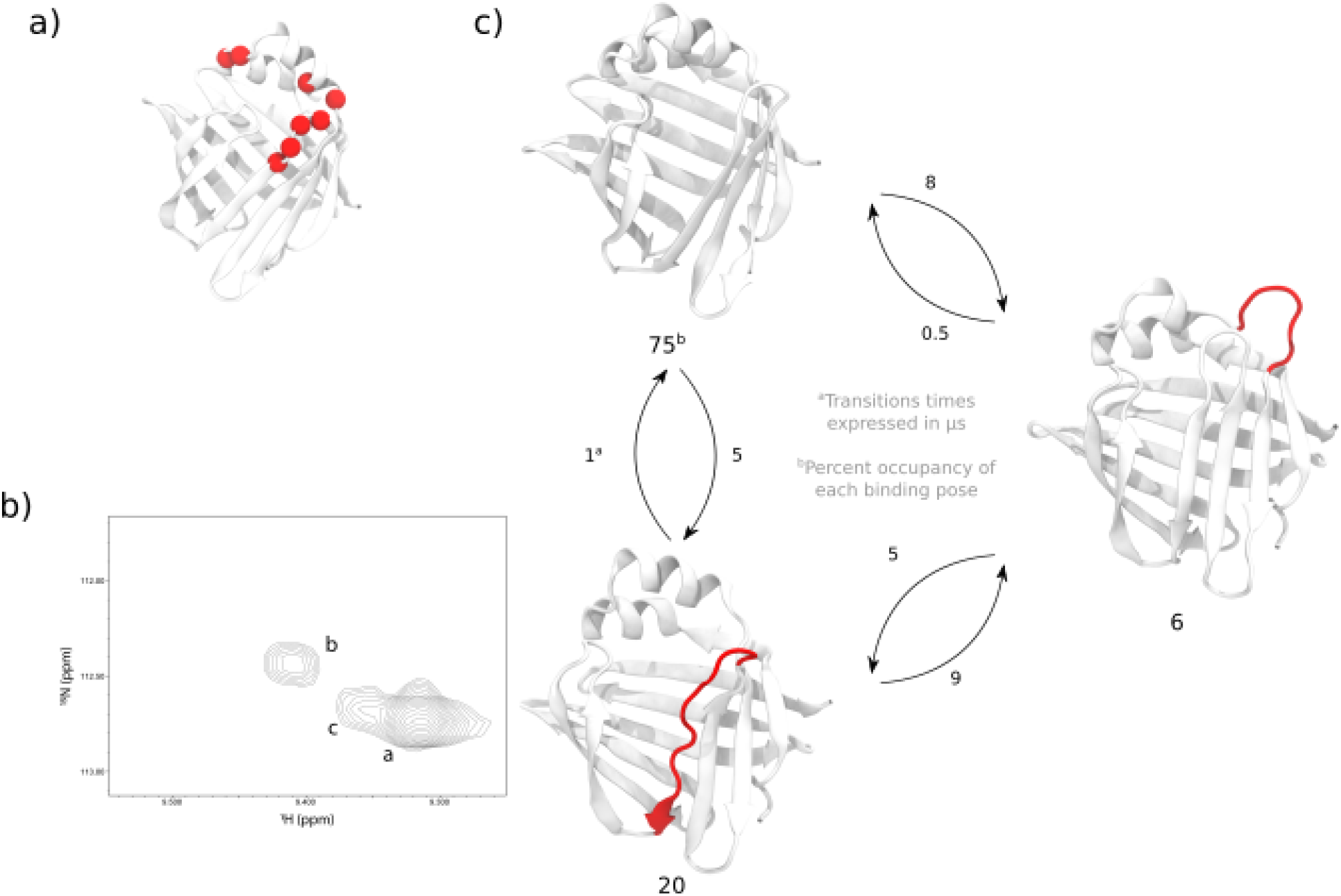
The portal region of apo FABP7 is highly dynamic and ligand binding leads to reduced dynamics. (a) Location of residues that are assigned multiple HSQC peaks. (b) Multiple peaks assigned to T60. (c) Hidden Markov model-generated kinetic network of metastable states for apo FABP7. Red colour of protein indicates unfolding in that region.

Using molecular dynamics simulations, we sought to understand the protein dynamics that give rise to the multiple HSQC peaks. Briefly, we use hidden Markov models paired with molecular dynamics data to characterize the long-timescale dynamics of FABP7. From this analysis, we identify alternative states from the native fold characterized by the partial unfolding of H3 or βD (Fig. 4c).

The H3-unfolded state is associated with less persistent backbone hydrogen bonds between residue pairs V32/T36 and T29/G33 compared to the native fold. Similarly, for the partially unfolded βD state, T54 and T60 have diminished hydrogen bonding (Figs. S9 and S10). These partially unfolded states are associated with flexible βCD/βEF turns and alpha helices (Fig. S11). The multiple 2D ^1^H-^15^N HSQC peaks observed for T36 and T60 may arise due to states with partial unfolding to H3 or βD.

Despite the correlation between NMR and simulation, the minor states observed in the simulations do not fully explain the NMR data. Multiple peaks are observed in the turn between H2 and H3, yet the simulations do not predict significant structural changes in this region and only predict unfolding in the last turn of H3 (residues 29 to 33). Moreover, the existence of multiple 2D ^1^H-^15^N HSQC peaks for a residue suggests that there are states that transition on the high µs to ms timescale, while the HMM predicts low µs transition times. It is unlikely that the predicted states fully explain the timescales captured by the HSQC spectra.

Unfolding to H3 and the gap region has also been observed for other apo FABPs. Extensive NMR and simulation studies performed on intestinal-FABP (FABP2) (32, 34–37) and epidermal-FABP (FABP5) (33, 38–40) reveal that, like FABP7, these proteins also undergo H3 and gap region unfolding. For FABP2, ^15^N relaxation dispersion (RD) and chemical shift saturation transfer (CEST) NMR experiments reveal minor populations with structural changes to H3 and gap region residues (34). X-ray crystallography structures of FABP5 suggest higher H3 flexibility (38), while data from hydrogen-deuterium exchange experiments (34–36) on both proteins show lower protection factors and increased flexibility for residues in H3.

Simulations of FABP2 and isolated secondary structure motifs reveal states with partially unfolded H3 (35), which is consistent with reduced FABP2 HDX protection factor data and our predicted dynamics of FABP7. Equilibrium simulations on apo FABP2 revealed an intermediate along the unfolding pathway, while full unfolding of H3 was not observed due to insufficient sampling (35). Nevertheless, metadynamics simulations captured minima corresponding to the native fold, partially unfolded H3, and a fully unfolded H3.

Combining our data with the apo structural dynamics of FABP2 and FABP5, we propose that the full unfolding of H3 or βD leads to the multiple peaks in HSQC experiments. Our predicted partially-unfolded states are likely intermediates along an unfolding pathway. The dynamics sampled by our simulations are not long enough to capture the entire H3- or βD-unfolding pathway, which likely occurs on longer timescales.

Taken together, we provide support for the presence of an H3- and βD-unfolded states that may be ubiquitous across the FABP family. For FABP2 (32, 34–37) and FABP5 (38–40), the unfolding of H3 occurs on a similar timescale to ligand binding, which may indicate that H3-unfolding is an essential step along the binding pathway (34). Regardless, these minor states may be necessary for the other biological activities of FABP7, namely in driving membrane interactions or nuclear localization.

### 4. Ligand binding stabilizes the portal region of FABP7

Our 2D ^1^H-^15^N HSQC spectra reveal several differences between apo- and holo-FABP7 (Figs. S12-S15). Regardless of the fatty acid bound, the largest chemical shift perturbations between apo- and holo-FABP7 are located at the βCD loop ends (T54, N59, and T60) and H3 (G33 and T36; Figs. 5 and S11). Notably, the residues with the largest chemical shift perturbations are identical to or located near residues with multiple peaks in the apo state, including G33 and T60. Furthermore, the saturated holo HSQC spectra no longer show multiple backbone peaks for these portal region residues indicating that ligand binding has a large effect on the protein heterogeneity or dynamics within these regions (Fig. S8). Although the multiple peaks observed for the apo-state may have moved from slow to fast exchange upon ligand binding, ligand binding stabilizes the portal region towards a single primary conformational macrostate. This supports the idea that these residues’ conformational state may signal ligand binding across the FABP family.

**Figure 5.**
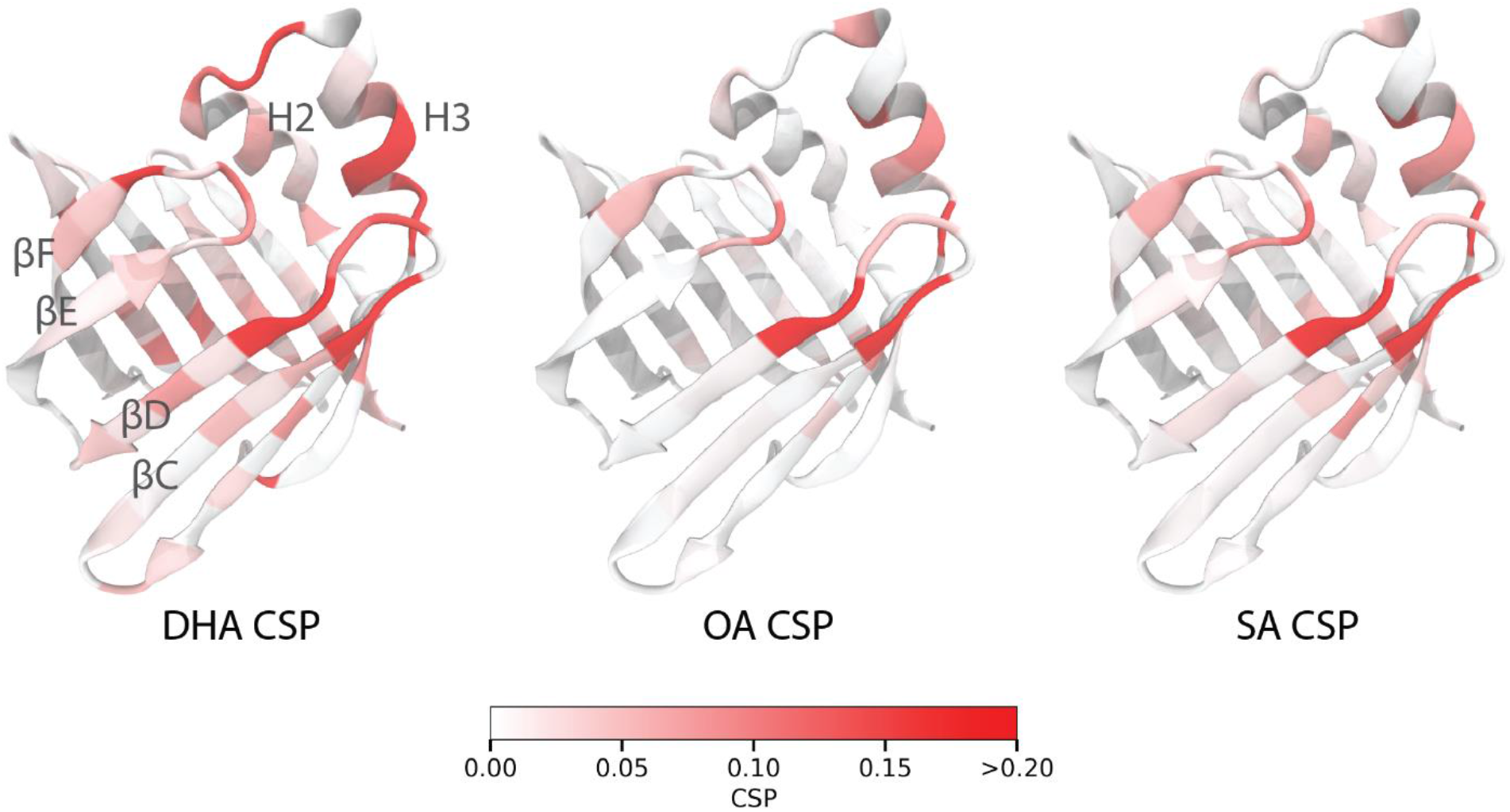
Fatty acid binding leads to perturbations in portal region residues. (a) CSPs between apo and DHA, OA and SA bound spectra are mapped onto the structure of FABP7. Largest CSP is observed with portal region residues for all FAs. Binding of DHA show more residues in H2-H3 and βCD/βEF turns with greater CSPs. Absolute CSP bar plots for each fatty acid can be found in Fig. S15.

To analyze how ligand binding changes FABP7 protein dynamics, we utilized time-independent coordinate analysis (TICA). Like principal component analysis (PCA), TICA is a dimensionality reduction method that linearly transforms input features. Unlike PCA, which finds coordinates that maximizes the input features’ variance, TICA finds coordinates that maximize the input features’ autocorrelation, making it a suitable method for analyzing slow dynamics.

For wild-type/apo FABP7, we inputted key protein-protein distances as features for TICA and plotted the first two components (Table S1-S2 and upper left panel in Fig. 6). These components are orthogonal coordinates corresponding to the slowest evolving protein dynamics, which qualitatively map to gap unfolding (primarily βD; y-axis) and H3 unfolding (x-axis) for our system (Fig. S9). The wild-type/apo simulations (top left panel of Fig. 6) show a distinct basin corresponding to the native fold. The native fold is separated by a small barrier from gap-unfolded structures, while structurally diverse conformations with low populations that involve the unfolding of H3 are also present. These alternative minor states are predicted to be associated with the portal region residues with multiple HSQC peaks in the apo spectra.

**Figure 6.**
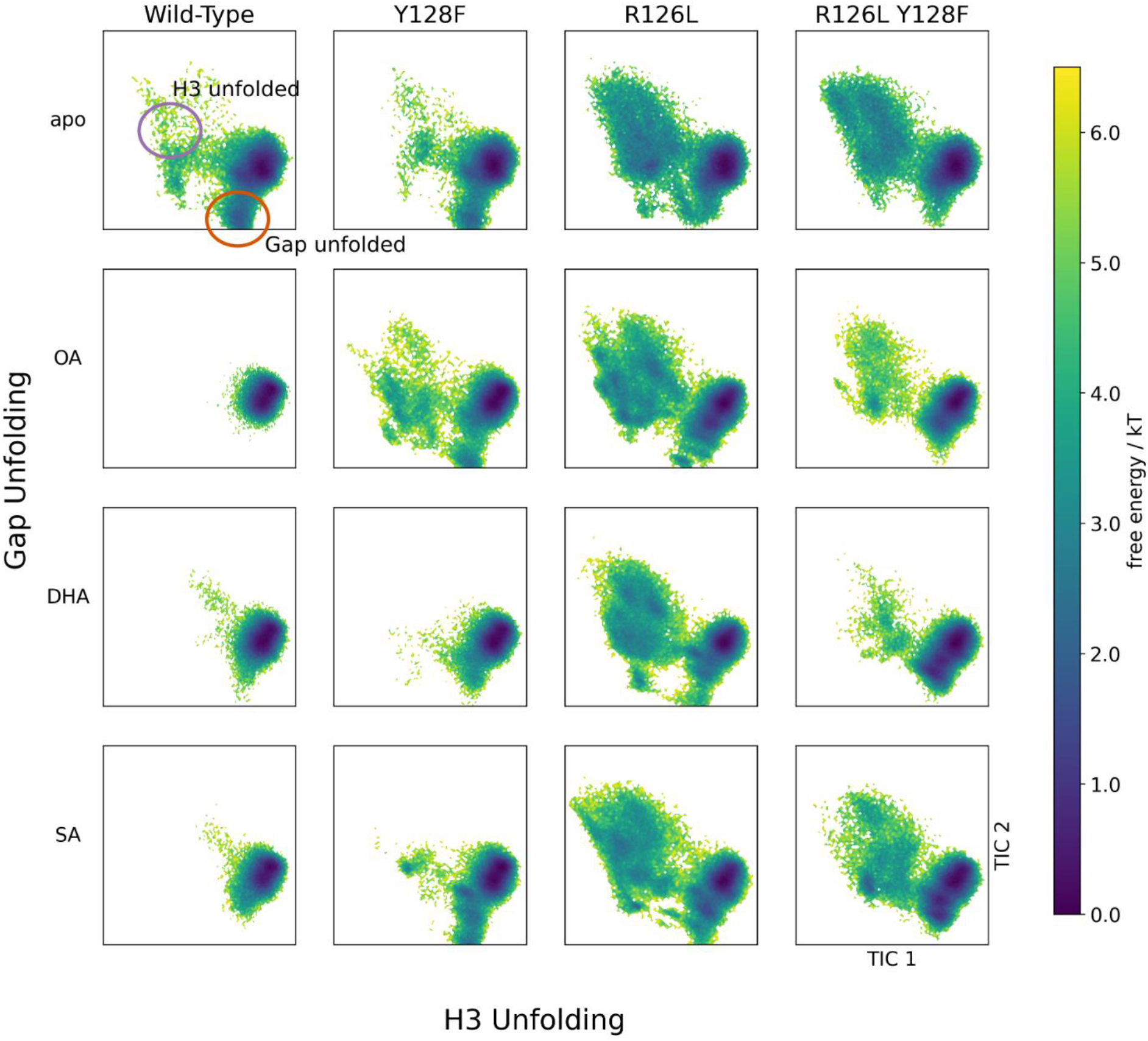
Ligand binding stabilizes the native fold of FABP7, while mutations to binding residues lead to unfolded H3 and gap region states. Each panel shows a system (labelled by row and column) projected onto the slowest TICA components of wild-type apo FABP7, which corresponds to features (listed in Table S1) that comprise gap unfolding (y axis) and H3-unfolding (x axis).

We projected the ligand-bound simulations onto the slowest evolving apo FABP7 TICA components. For the wild-type protein, the native fold of FABP7 is stabilized regardless of the ligand-bound (left column of Fig. 6). For wild-type holo-FABP7, we observe a disappearance of the partially unfolded gap and H3 structures present for the apo state.

The increased stability of the native fold is driven by tighter intra-protein interactions. Contact map occupancies show that contacts between βCD–H3 and βCD-βEF become more extensive upon ligand binding (Fig. 7). In particular, F57 tightly associates with T29, V32, and G33, while several contacts are more persistent between K58/T60 and the βEF turn. Interestingly, we also observe a decrease in protein fluorescence anisotropy upon ligand binding (Fig. S7). We interpret this as being associated with the closed compact protein tumbling faster in solution based on our structural data. The reduced conformational heterogeneity in the portal region upon ligand binding can explain why residues (G33, T36, T54, F57, N59, and T60) with multiple peaks in the apo HSQC spectra converge to single peaks in the holo HSQC spectra.

**Figure 7.**
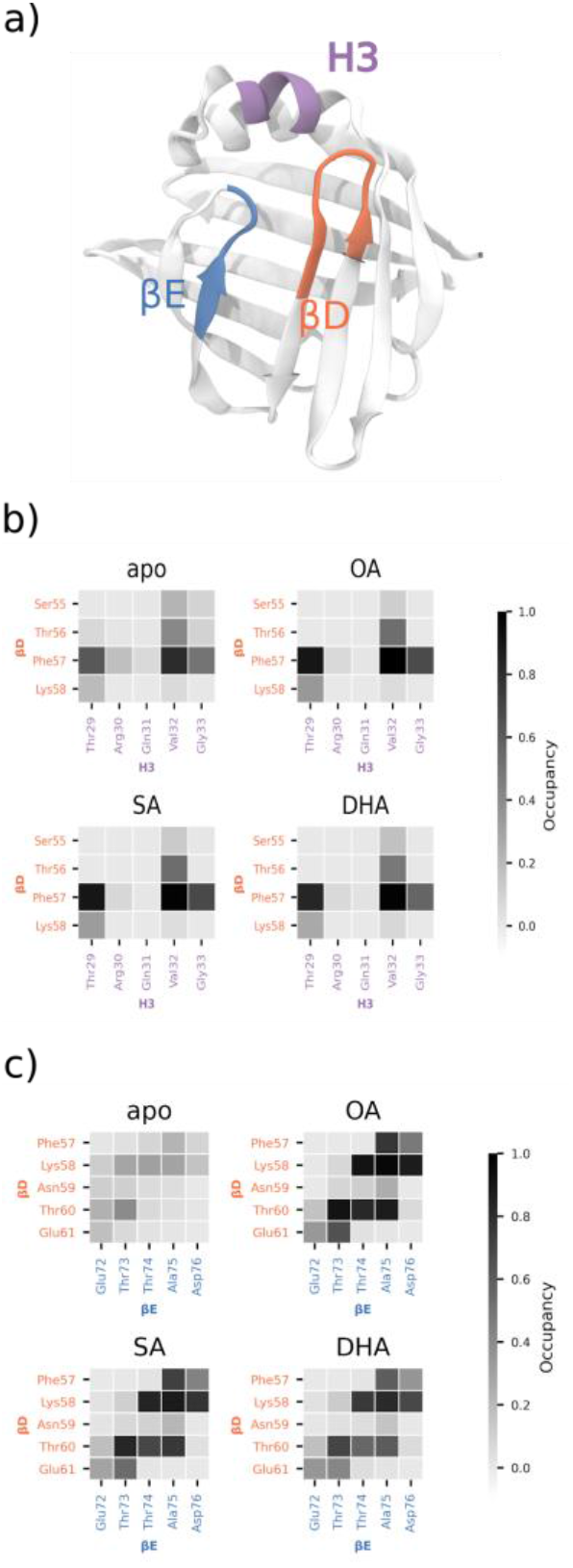
Ligand binding tightens the contacts between portal region residues. (a) colour scheme. (b and c) Shaded squares indicate contact occupancy (<4.5 Å separating closest two heavy atoms) between two residues in (b) βD and H3 or (c) βD and H3.

Our combined experimental and theoretical data suggests that the bound ligand stabilizes the portal region and prevents the unfolding of H3 and the gap region. Similar decreased protein conformational plasticity upon ligand binding has been identified for other FABP family members, including I-BABP (39) and H-FABP (33).

### 5. Changes to the dynamics of H3 may be key to understanding fatty acid-dependent biological outcomes

Despite only small differences in binding affinities between the ligands at 37°C, NMR data and molecular dynamics simulations reveal that each fatty acid has a unique effect on the conformation of FABP7.

Compared to apo FABP7, OA-bound FABP7 has the smallest chemical shift perturbations (Figs. 5 and S15). SA binding results in similar perturbations to portal region residues as OA but with a larger perturbation to N59. Intriguingly, DHA results in significant perturbations to residues in H3 and the βCD/βEF turns (Fig. 5). This includes a perturbation to NLS residue Q31 that is not observed when the other ligands are bound. Therefore, we propose that DHA binding permits greater H3 unfolding, which affects the structure and dynamics of the NLS uniquely. This change to the protein structure and dynamics may lead to the localization of the FABP7-DHA complex to the nucleus.

Our simulation data correlates with the significant perturbations observed by NMR for DHA binding to FABP7 compared to the other ligands. Specifically, DHA binding leads to increased flexibility and weaker contacts among gap region residues compared to OA or SA (Fig. 7). However, only modest differences are observed in H3 among the ligands. It is possible that DHA binding perturbs the dynamics of H3 on timescales that we have not sampled with our simulations, as was the case for apo FABP7.

Our data reveal that DHA binding leads to changes to H3 and βCD/βEF turns of FABP7 that do not occur upon OA or SA binding. Therefore, we propose that DHA-induced changes to the portal region’s conformational dynamics contribute to the nuclear localization of FABP7. This is consistent with our binding assays that show little difference in binding affinity at 37ºC for any of the three ligands.

### 6. Binding pose heterogeneity explains changes to protein dynamics and mutation-resistant binding

While we observe that DHA binding to FABP7 uniquely perturbs the structure of H3, it is unclear why mutation of key binding residues have little effect on binding of the three fatty acids studied (Fig. 3). Indeed, previous studies indicate that mutation of R126 and Y128 to alanine reduces FABP7’s binding affinity for fatty acids (5). Even though our mutations are isosteric (R126L and Y128F) and may have less of an impact than mutation to alanine, we hypothesized that the effect of binding affinity would be larger than what we observed. Using molecular dynamics and hidden Markov models, we analyze how the ligands bind to wild-type and mutant FABP7 to understand why R126L and/or Y128F mutants maintain tight binding to fatty acids.

We observe three unique binding conformations in the FABP7 and fatty acid complex. The two most populated conformations have a hydrogen bond between the ligand and both R126 and Y128 (top and bottom left Fig. 8). The poses differ based on the tail conformation. The most populated conformation orients the lipid tail towards βD (top) and is most similar to the bound conformation adopted by OA and DHA in X-ray crystal structures (Fig. S16) (16). Specifically, the OA adopts a U-shape conformation, while the DHA tail adopts a helix-like conformation, and interactions with F16 and F104 stabilize both tail conformations. The binding pose of DHA does not precisely match the crystal structure with the tail not looping across the gap region, which may be due to the flexibility and heterogeneity of the lipid tail. The flexible lipid tail is consistent with NMR data that suggests the C12-C18 carbons of the FABP3-bound fatty acids become disordered at room temperature (41).

**Figure 8.**
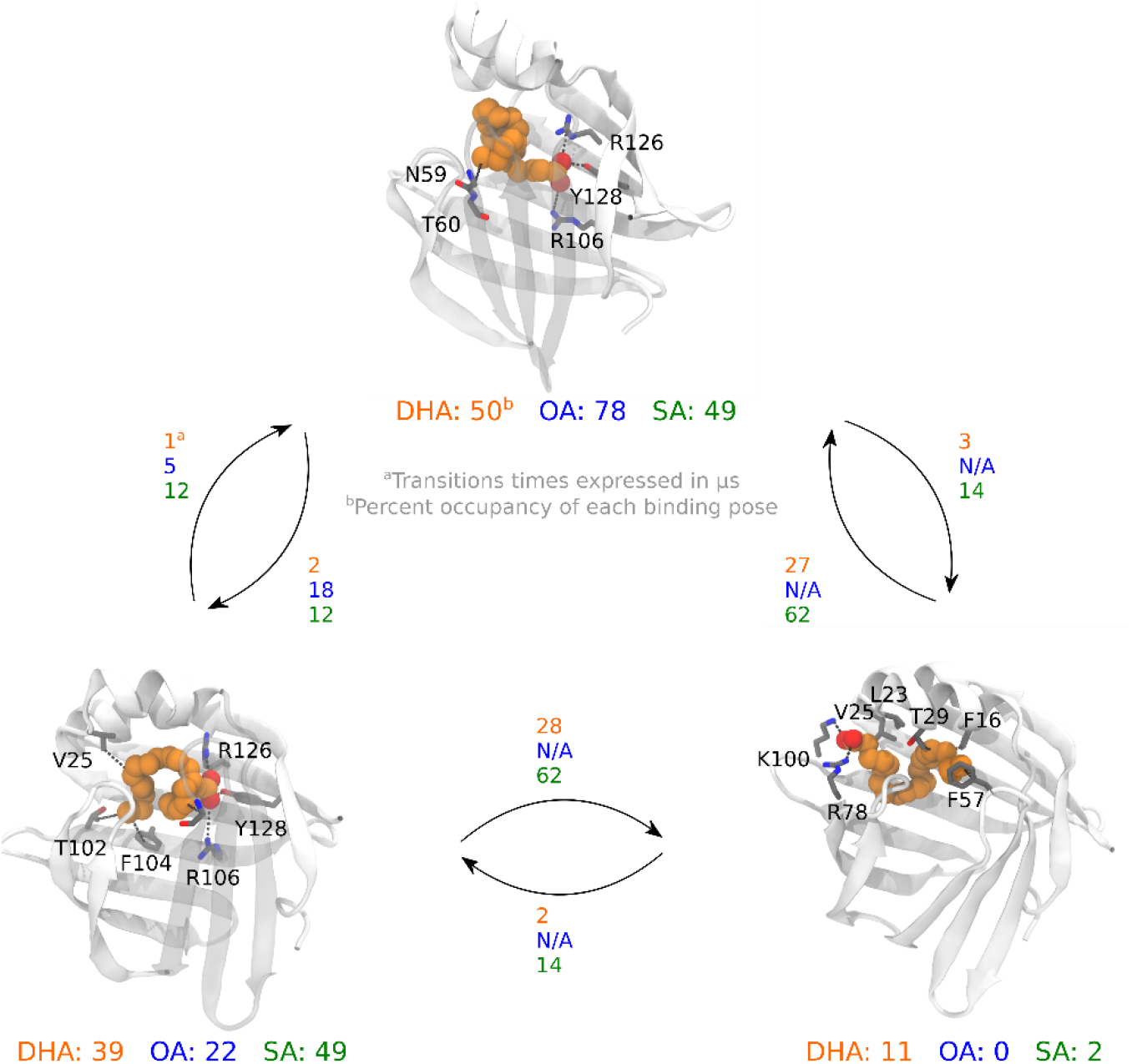
Each fatty acid exhibits a dynamic lipid tail when bound and an alternative carboxylate binding site exists for DHA and SA. Hidden Markov model-generated kinetic network of ligand binding poses for DHA, OA, and SA. For clarity, only the pose of DHA is shown in orange and key binding residues in gray.

A third binding pose has the ligand carboxylate interacting with R78 and K100, with the tail extending toward H3 (right, Fig. 8). This conformation is more probable with DHA (11% occupancy) than SA (2% occupancy) and is not observed for OA. DHA’s higher occupancy may be due to its tail length and flexibility and stabilized by hydrophobic interactions with T29, L23, V25, F16, and F57. The location of these interactions is near H3 and help explain why larger significant HSQC CSP occur upon binding DHA compared to the other fatty acids.

To understand why R126L and/or Y128F mutants had little impact on the binding affinity of FABP7 towards OA, DHA, or SA (Fig. 3), we examined the ligand-protein dynamics using hidden Markov models. We summarized the data by analyzing 1000 HMM-generated structures from each unique system with PCA (Fig. 9). Using HMMs to generate structures leads to greater equilibrium sampling by reducing the structural bias associated with the docked starting structures. Subsequently, we combined the structures from all systems, calculated 9 distances that correspond to different ligand-binding sites, and performed PCA on the results to project each system onto a consistent coordinate system. The structures are colored according to their varying conformations (Fig. 9).

**Figure 9.**
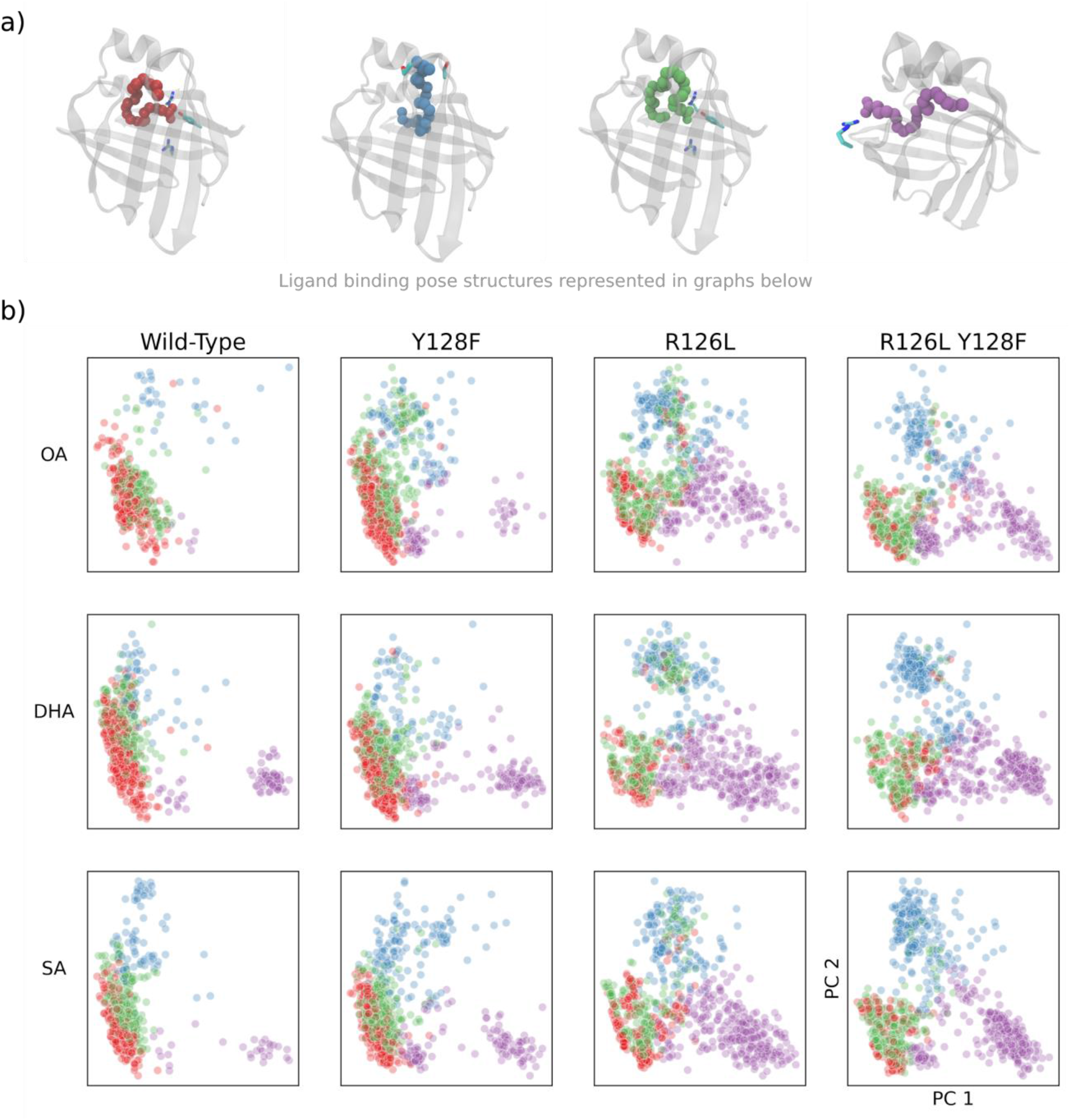
R126L and/or Y128F mutations lead to fatty acids sampling alternative conformations more frequently. PCA was used to combine coordinates corresponding to different ligand binding sites for 1000 HMM-generated structures from each system. The structures from each system are projected onto that coordinate system to generate each panel in the figure corresponding to the ligand bound and the FABP7 phenotype.

For the wild-type systems, PCA analysis identifies a fourth binding conformation with the carboxylate hydrogen bonding to T36 and T55. Based on our HMMs, this pose is not kinetically distinct from other poses and rapidly transitions to structures where the ligand interacts with R126 and Y128. Despite this new binding conformer, the observed ligand binding poses usually interact with R126/Y128.

In general, the R126L and Y128F mutations increase the heterogeneity of the ligand binding poses adopted in the FABP7 binding pocket (Fig. 9). Each mutant increases the frequency of the T36/T55- and R78/K100-interacting binding poses being adopted, with the Y128F mutation having a smaller effect compared to R126L. The diversity within each major binding pose increases depending on the mutation. Specifically, with R126L mutation, there is a more significant overlap between the R126/Y128-interacting binding pose and more heterogeneity within the other poses.

For both apo and holo FABP7, the R126L mutation has a larger effect on the diversity of protein conformations than the Y128F mutation (Fig. 7). This is because R126 forms water-mediated hydrogen bonds with G33 and N34 backbones (Fig. S17), while these interactions are not present for the R126L mutant. Losing these hydrogen bonds destabilizes the secondary structure of H3, leading to a higher proportion of H3-unfolded structures, and suggesting that R126 may also regulate the conformation of H3 in addition to its ligand binding role. The water-mediated hydrogen bond between R126 and H3 is also present in FABP3 (42).

The mutation-resistant binding we observe via MST (Fig. 3) is explained by the ability of FABP7 to accommodate fatty acids in several structurally distinct conformations, which permits binding in the absence of hydrogen bond donors at the R126 or Y128 positions (Fig. 9). Even for wild-type FABP7, there is still considerable structural diversity in the binding poses observed. More broadly, the ability for FABP7 to bind fatty acids in several conformations may be a factor in its promiscuous binding to structurally diverse ligands, including endocannabinoids (10, 11), phytocannabinoids (12), and dyes (13) that may locate to different regions of the binding pocket. To the best of our knowledge, this is the first study to demonstrate that an FABP exhibits mutation-resistant binding, and we also provide evidence that the existence of multiple redundant binding conformations leads to this phenomenon.

## Conclusions

By using a variety of biophysical techniques, we provide fundamental insight into how FABP7 binds fatty acids and how each fatty acid uniquely affects the protein structure. We establish that no fatty acid binding preference exists at biologically-relevant temperatures. Our simulations and NMR data reveal that the portal region of apo FABP7 is highly dynamic, and ligand binding leads to unique changes to these dynamics depending on the fatty acid bound. Notably, when DHA is bound, the NLS-containing H3 is uniquely perturbed, which provides clues as to why the FABP7-DHA complex is preferentially relocated to the nucleus. We also establish that FABP7’s fatty acid binding affinity is resistant to mutation of key binding site residues, and that this resistance is conferred by FABP7’s ability to bind fatty acids in several unique binding poses. Overall, we provide comprehensive insights into how FABP7 interacts with key ligands and how these interactions may govern FABP7’s behavior within the cell.

## Supporting information

supplementary methods, figures, tables, and references

## Author Contributions

I.B. conceptualized the study, performed all NMR experiments, analyzed NMR data, and wrote and edited the article. S.L. conceptualized the study, performed all computational modelling and analysis, and wrote and edited the article. M.R.Y. conceptualized the study, secured funding, performed all ANS, MST, and melting temperature experiments, analyzed the corresponding data, and wrote and edited the article. T.S. analyzed experimental data and edited the article. H.I. analyzed NMR data and edited the article. H.J.V. analyzed NMR data and edited the article. J.L.M. conceptualized the study, secured funding, supervised the study, analyzed experimental and computational data, and wrote and edited the article.

## Acknowledgements

The authors thank the Natural Sciences and Engineering Research Council of Canada (NSERC) for financial support through its ENGAGE program (ENGAGE: EPG-488413-15, ENGAGE Plus: EPG2-499238-16 and Alcohol Countermeasures Corp, in particular Maciej Goledzinowski and Felix J.E. Comeau, for their overall support for this work. M.R.Y. thanks NSERC for support through an NSERC postdoctoral fellowship and S.L. thanks the Canada First Research Excellence Fund for his postdoctoral fellowship. The authors also thank the Canada Research Chairs and Discovery Grant programs for their support of J.L.M. The authors are grateful for the computer resources provided by Compute Canada.

