## supplementary methods, figures, tables, and references for "Brain fatty acid binding protein exhibits non-preferential and mutation-resistant binding towards fatty acids"

Iulia Bodnariuc*, Stefan Lenz*, Margaret Renaud-Young*, Tanille Shandro, Hiroaki Ishida, Hans J. Vogel, Justin L. MacCallum

* equal first author

#### Table of Contents

Full Experimental and Computational Methods ........ S3
Supplementary Tables........ S5
**Table S1. Inverse distance features used for TICA analysis of apo FABP7.**........ S5
**Table S2. Inverse distance features used for TICA analysis of FABP7 bound to DHA.** ........ S6
Supplementary Figures ........ S7
**Figure S1**. Docked starting structures for simulations of FABP7 bound to a) OA b) DHA and c) SA. ........ S7
**Figure S2.** a) Implied timescales and b) Chapman-Kolmogorov plots for HMM built using simulations of FABP7 bound to DHA. ........ S8
**Figure S3.** MST binding curves of wtFABP7 for each fatty acid at 25, 29, 31, and 37 ºC. This data was used for the Van't Hoff analysis discussed in Figure 2. ........ S9
**Figure S4**. Binding curves for OA (blue), DHA (orange), and SA (green) generated from MST experiments for Dmut (double mutant, R126L/Y128F) and R126L FABP7 at 25, 31, and 37 ºC... S10
**Figure S5.** Van't Hoff Analysis of MST binding data for wt, R126L, and R16L/Y128F FABP7 show fatty acid binding is temperature dependent. ........ S11
**Figure S6.** a) The ANS-protein binding curves used to determine ANS dissociation constant b) ANS displacement assay curves, concentration of ANS is set to 1μM, the halfway point of this curve corresponds to the $EC_{50}$ of OA (blue), DHA (orange) and SA (green) c) shows the table of vaues measured for each curve and the associated calculated $K_i$ value from $K_i = EC_{50}/(1+[ANS]/K_{D,\,ANS})$ ........ S12
**Figure S7.** Fluorescence anisotropy for increasing concentrations of OA and DHA bound to FABP7. Upon Ligand binding we see a decrease in fluorescence anisotropy. The dissociation constants determined for OA (blue), DHA (orange), and SA (green) are shown in the bottom left of the graph. ........ S13
**Figure S8**. Residues which show multiple peaks in the $^{1}H$-$^{15}N$ HSQC of the apo-wtFABP7. Titration of ligand shows all three fatty acids (DHA, OA, SA) are all in slow exchange and the multiple peaks observed collapse to a single peak upon ligand saturation. ........ S14

**Figure S9.** H3 and βD unfolded states are associated with reduced intra protein hydrogen bonds. Native (black) and unfolded state (red) occupancies are labelled next to dashed lines denoting hydrogen bonds........ S15
**Figure S10.** Unfolding of H3 and gap region result from loosening of contacts between the βCD turn and H3 or βEF turn. Input contact features (represented as purple cylinders) that elongate upon unfolding for the a) H3-unfolded state and b) gap-unfolded state. The unfolding of these regions is primarily driven by the presence or absence of contacts between the βCD turn and H3 or the βEF turn. ........ S16
**Figure S11.** rmsf differences plots reveal that the βCD and βEF turns are more flexible for SA and DHA compared to OA with DHA having the largest difference. Absolute Cα rmsf for a) apo FABP7 and b) FABP7-OA. Cα rmsf difference between OA and c) DHA and d) SA. ........ S17
**Figure S12.** $^{1}$H-$^{15}$N HSQC showing 50μM apo-wt FABP7 (gray) and 50μM holo-wtFABP7 saturated with 300μM DHA (yellow). Peaks of importance are labeled on spectra. Assignment was imposed with literature values (13). ........ S18
**Figure S13.** $^{1}$H-$^{15}$N HSQC showing 50μM apo-wt FABP7 (gray) and 50μM holo-wtFABP7 saturated with 250μM OA (blue). Peaks of importance are labeled on spectra. Assignment was imposed with literature values (13)........ S19
**Figure S14**. $^{1}$H-$^{15}$N HSQC showing 50μM apo-wt FABP7 (gray) and 50μM holo-wtFABP7 saturated with 300μM SA (green). Peaks of importance are labeled on spectra. Assignment was completed by peak following and comparison to literature values (13). ........ S20
**Figure S15.** CSP for each residue associated with fatty acid binding of a) DHA, b) OA, and c) SA. ........ S21
**Figure S16.** Superposition of HMM-sampled structures from most populous state for a,c) OA and b,d) DHA onto crystal structures of FABP7 bound to OA (PDB:1fe3) or DHA (PDB:1fdq). For a) and b) 10 structures were randomly sampled to provide a qualitative estimate of the ligand binding pose heterogeneity, while c) and d) show the binding pocket for a single HMM-sampled structure. ..... S22
**Figure S17.** Water-mediated hydrogen bonds (water oxygen atoms are represented as red spheres) between R126 and G33, N34, and T36 are lost upon mutation of R126 leading to greater H3 instability. ........ S23
References ........ S24

### Full Experimental and Computational Methods

#### Experimental

*ANS Displacement Assay*

Binding interaction measurements were performed in the same manner for FABP7 wild-type and mutant proteins. 1-anilino-8- naphthalene sulfonate (ANS) signal intensity increases 40-fold when in the hydrophobic environment of the protein binding pocket. ANS binding affinity was determined by performing a dilution series of FABP7 with a constant value of ANS (1 μM). An inhibitory constant ($K_i$) for FABP7 (wildtype and mutants) was determined by combining 1 μM of FABP7, 5 μM ANS and dilution series of OA, DHA or SA. Samples were incubated for greater than 30 minutes at 25 °C, then transferred to a 384 well black plate (Greiner). The Ki value was calculated using:

$$K_i = {}^{EC_{50}}\Big/_{(1 + {}^{[ANS]}\big/_{K_{D\,(ANS)}}}$$

*Microscale Thermophoresis (MST)*

Microscale Thermophoresis measurements were carried out using Monolith NT.115Pico instrument. Measurements were carried out in MST-buffer (20 mM phosphate, 120 mM NaCl, at pH 7.4, in 0.005% Tween-20) using premium capillaries (Nanotemper). Proteins were labeled with RED-tris-NHS dye following supplier protocol. Measurements were taken with instrument set to high MST power, and 20-30% LED power. A solution of 20 nM FABP7-NHS-RED-Tris labeled was combined with fatty acid dilution series. Samples were incubated for 30 minutes at measurement temperature before loading samples into MST capillaries. Experiemnts were completed with 3 replicate dilution series. Analysis of MST traces was completed using MO.Affinity Analysis v2.3 (Nanotemper) (1).

#### Computational

*Molecular Dynamics.* The crystal structure of FABP7 bound to antinociceptive SBFI-26 (PDB ID: 5URA) (2) was used as the starting structure for our models. To simulate apo FABP7, the inhibitor was removed, while hydrogen atoms were initially assigned with H++(3) and adjusted to maintain hydrogen-bonding networks within the protein. The systems were solvated with a 10 Å octahedral water box (~5000 waters, side length ~61 Å) and neutralized with sodium ions using the tLEaP utility from AmberTools (4).

The OpenMM (5) python library was utilized to run all molecular dynamics simulations. The Amber ff14SB forcefield (6) was used for all protein atoms. Each simulation was minimized/equilibrated using the following protocol. The system was subjected to three 500 cycle rounds of limited-memory Broyden-Fletcher-Goldfarb-Shanno minimization applied to: (1) water and ions (100 kcal $mol^{-1}$ $Å^{-2}$ restraint on atoms excluded from minimization); (2) protein backbone (25 kcal $mol^{-1}$ $Å^{-2}$ restraint); and 3) full system. Subsequently, the system was gradually heated to 310 K over 250 ps with a 25 kcal $mol^{-1}$ $Å^{-2}$ restraint placed on the complex and subjected to Langevin dynamics (1 fs timestep and 1 $ps^{-1}$ collision frequency). The restraint was released over 250 ps at 310 K under NPT conditions (1 bar) controlled by a Monte Carlo barostat. Constraints were applied to bonds involving hydrogen and the system was allowed to equilibrate for 500 ns using a 2 fs timestep. For all simulations, particle mesh Ewald (PME) was applied to electrostatic interactions and all non-bonded interactions were cutoff at 10 Å

*Docking.* To obtain starting structures from an ensemble of open and closed apo FABP7 conformations, 300 evenly spaced structures were taken from a 1 μs simulation of apo FABP7. Subsequently, AutoDock Vina (7) was used to generate docked poses of each ligand docked to each of the 300 apo structures. Poses within 1.0 Autodock Vina scoring unit from the tightest binding pose were retained (usually around 10 poses per apo structure). The generated complexes were then clustered according to the position of the binding pocket residues and the bound ligand to 30 clusters using Ward's hierarchical agglomerative clustering algorithm using cpptraj (8). This procedure produced several unique binding poses for each fatty acid bound to open and closed FABP7 (Figure S1). The medioid of each cluster was used as starting structures for MD simulations. In total, at least 50 μs of simulation time was obtained per system for a total of 800 μs of sampling time. Snapshots were saved and analyzed every 0.5 ns.

*Ligand Parametrization.* Each ligand was optimized at the B3LYP/6-31G(d) level of theory using Gaussian 16 (Revision C.01) (9). Following optimization, each ligand was assigned GAFF (10) atom types and AM1-BCC charges using the Antechamber utility from AmberTools (4).

*Hidden Markov State Modeling.* Hidden Markov models were generated using the PyEMMA version 2.5.7 (11). The input MD coordinates were first analyzed with time-independent coordinate analysis (tICA) (12) to reduce dimensionality and isolate slowly evolving modes. Specifically, ~30-40 combined protein-protein or ligand-protein distances (Table S1-S2) were measured for each trajectory and tICA (tiCA lag time = 5 ns) was used to find the linear combination of these coordinates that result in the longest autocorrelation times. This method generally resulted in ~10 tICA components. These components were clustered using kmeans to 100 clusters using PyEMMA default settings.

The clusters were used as input to generate hidden markov models. Estimation of hidden markov models are initialized by first estimating a markov model that has been coarse grained into macrostates using Perron-cluster cluster analysis (PCCA++). Implied timescale plots and Chapman-Kolmogorov tests were used to select the number of hidden macrostates and shortest lag time that generated stable models (Figure S2). For each generated hidden markov model, this resulted in ~2-4 states with a lag time of 10-20 ns. From the hidden markov models, transition times and stationary distributions of each macrostate were calculated.

### Supplementary Tables

**Table S1. Inverse distance features used for TICA analysis of apo FABP7.**

| Residue 1 | Atom 1 | Residue 2 | Atom 2 |
|---|---|---|---|
| F57 | Cζ | Q31 | Cγ |
| F57 | Cζ | A75 | O |
| T53 | Cγ1 | T60 | Cγ1 |
| K58 | Nζ | T74 | O |
| K58 | Nζ | D77 | Oδ2 |
| D17 | Oδ1 | N34 | Nδ2 |
| M20 | Cε | N34 | Oδ1 |
| M20 | Cε | R30 | O |
| K21 | Cγ | R30 | NH1 |
| T29 | O | V32 | Cγ1 |
| R30 | Cγ | N34 | Nδ2 |
| V32 | O | T36 | N |
| T56 | Cγ1 | T36 | Cγ1 |
| F57 | Cζ | T29 | Cγ2 |
| F57 | Cβ | T36 | Cγ2 |
| F57 | Cγ | V32 | Cα |

**Table S2. Inverse distance features used for TICA analysis of FABP7 bound to DHA.**

| Residue 1 | Atom 1 | Residue 2 | Atom 2 |
|---|---|---|---|
| DHA | C1 | R78 | Cζ |
| DHA | C2 | T102 | Cγ2 |
| DHA | C3 | F104 | Cζ |
| DHA | C20 | V32 | Cβ |
| DHA | C12 | F16 | Cζ |
| DHA | C22 | F16 | Cζ |
| DHA | C17 | T53 | Cγ2 |
| DHA | C1 | W97 | Nε1 |
| DHA | C22 | T60 | Cγ2 |
| DHA | C22 | T53 | Cγ2 |
| DHA | C1 | Y128 | OH |
| DHA | C1 | R126 | Cζ |
| DHA | C22 | W97 | Cζ2 |
| DHA | C4 | F16 | Cζ |
| DHA | C22 | R78 | Cδ |
| DHA | C16 | V25 | Cβ |
| DHA | C22 | T102 | Cγ2 |
| DHA | C21 | F104 | Cγ |
| DHA | C1 | R106 | Cζ |
| DHA | C13 | F57 | Cζ |
| DHA | C6 | T53 | Cγ2 |
| DHA | C3 | T60 | Cγ2 |
| K58 | Nζ | D77 | Cγ |

#### Supplementary Figures

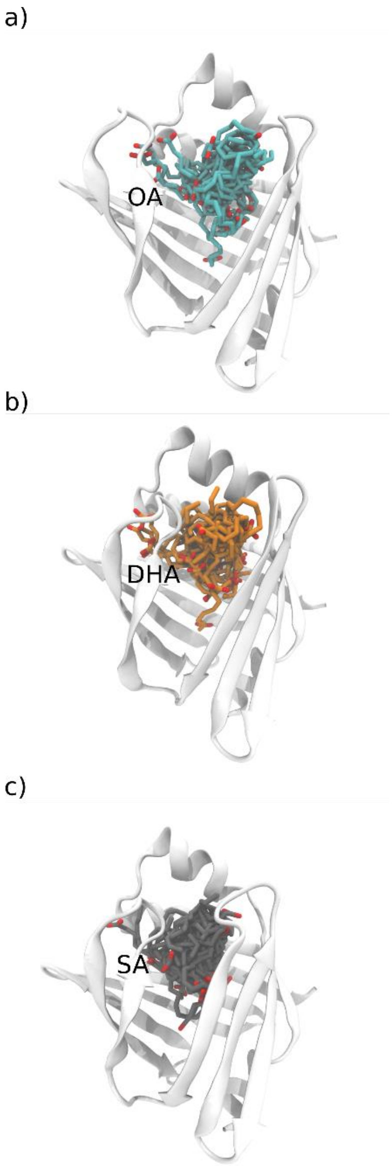


**Figure S1**. Docked starting structures for simulations of FABP7 bound to a) OA b) DHA and c) SA.

a)

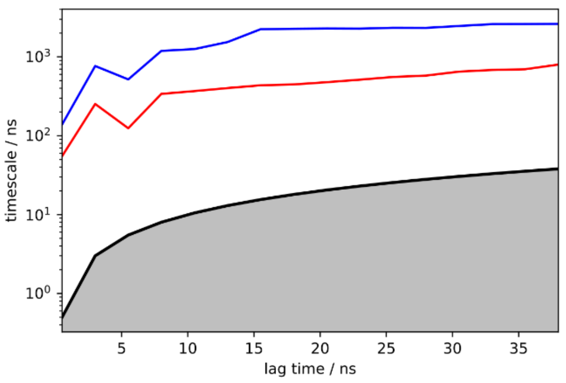


b)

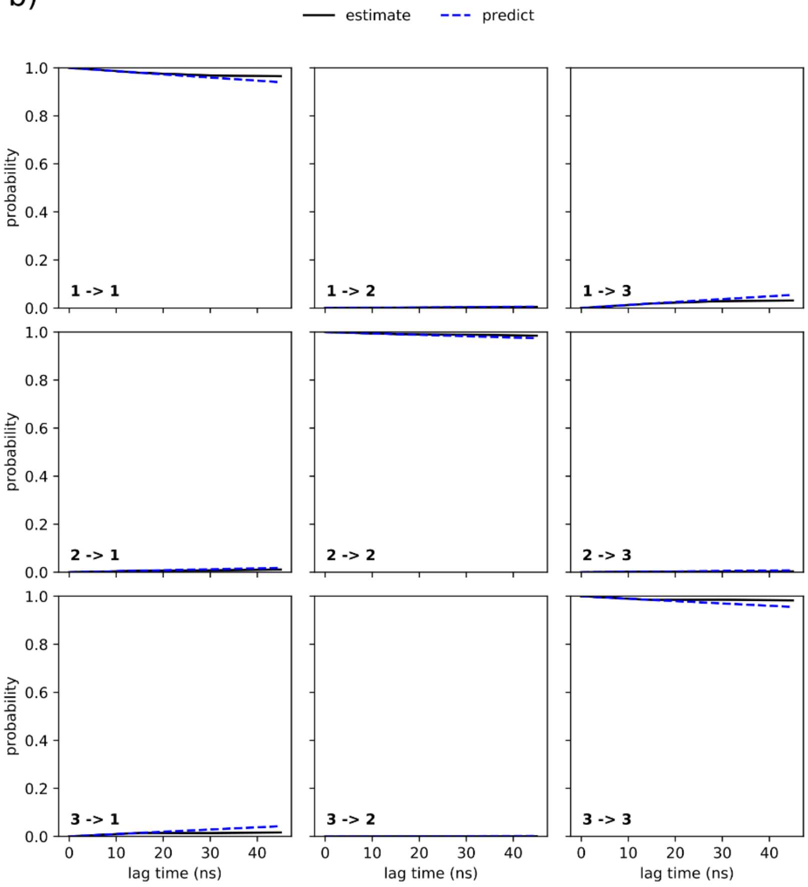


**Figure S2.** a) Implied timescales and b) Chapman-Kolmogorov plots for HMM built using simulations of FABP7 bound to DHA.

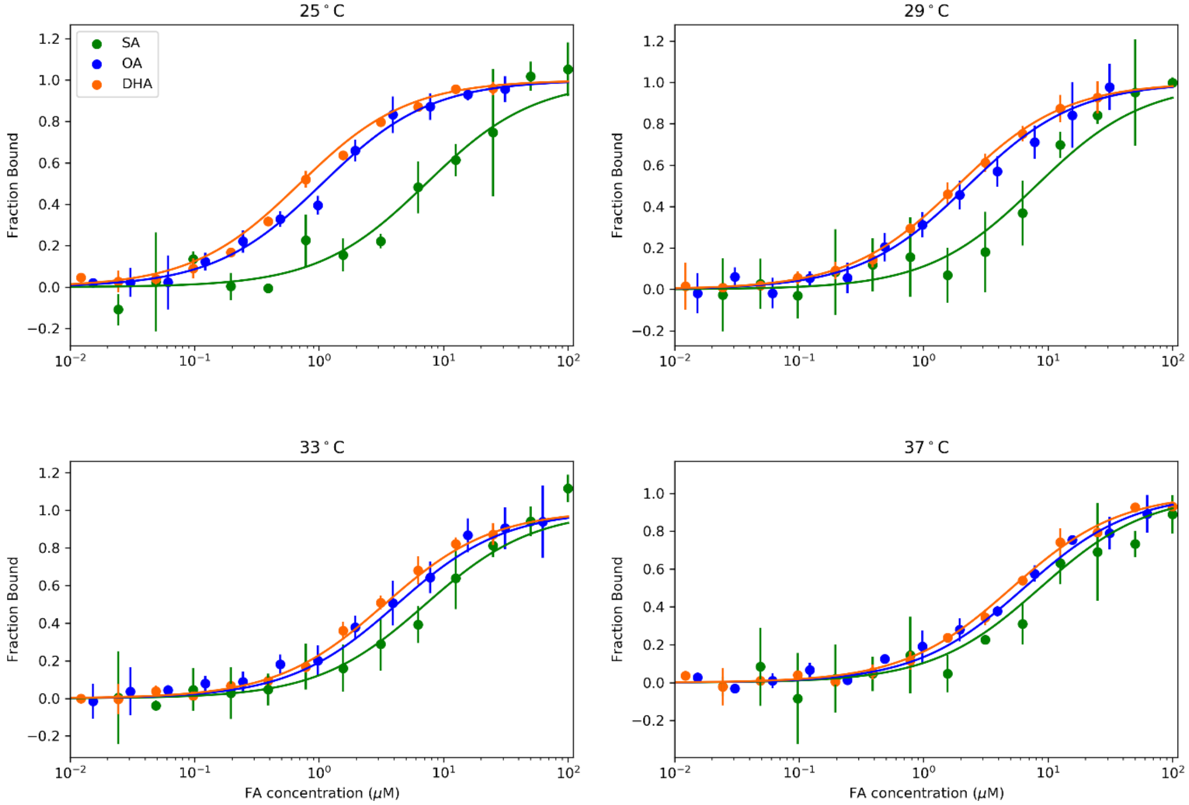


**Figure S3.** MST binding curves of wtFABP7 for each fatty acid at 25, 29, 31, and 37 ºC. This data was used for the Van't Hoff analysis discussed in Figure 2.

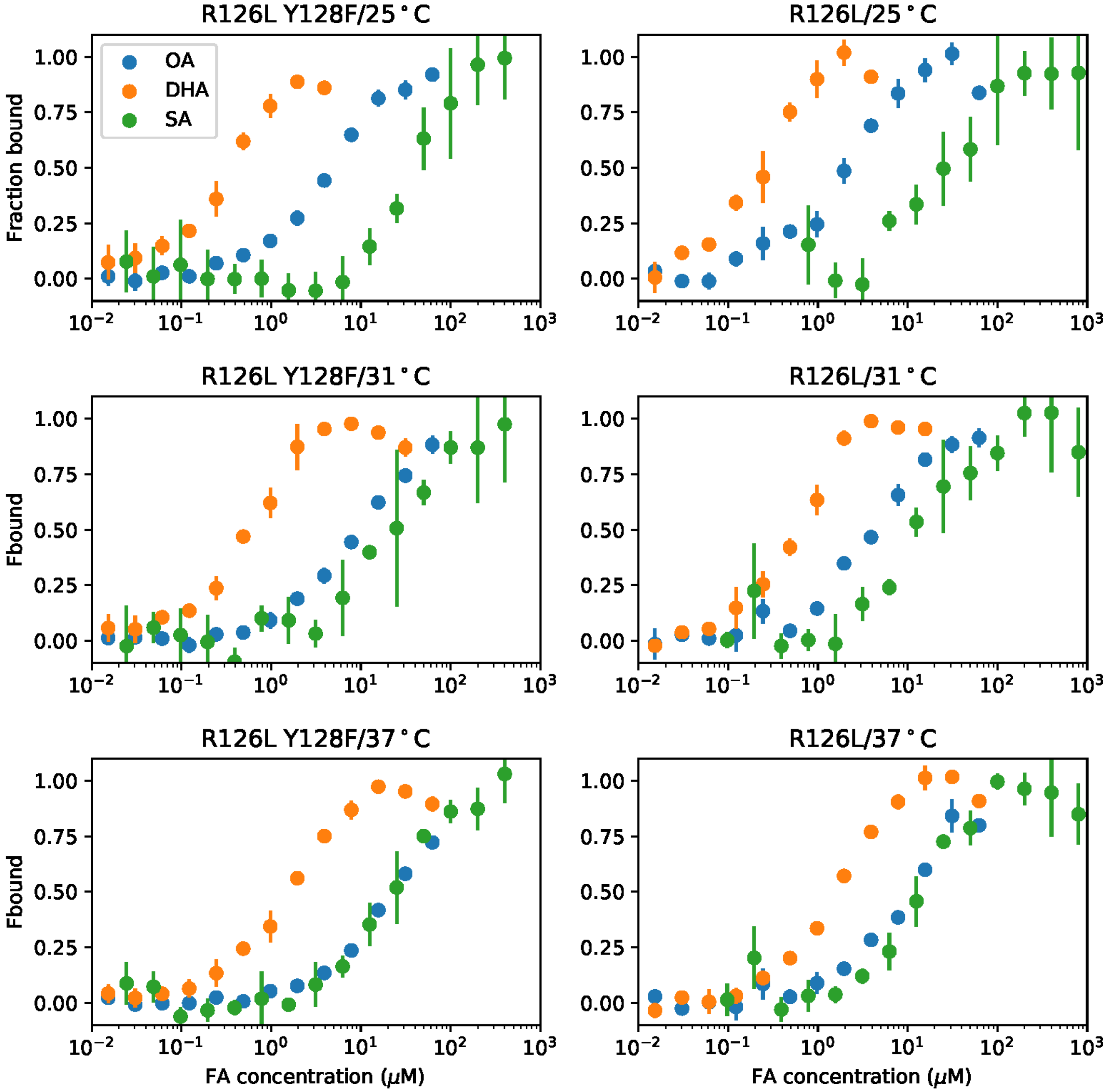


**Figure S4**. Binding curves for OA (blue), DHA (orange), and SA (green) generated from MST experiments for Dmut (double mutant, R126L/Y128F) and R126L FABP7 at 25, 31, and 37 ºC.

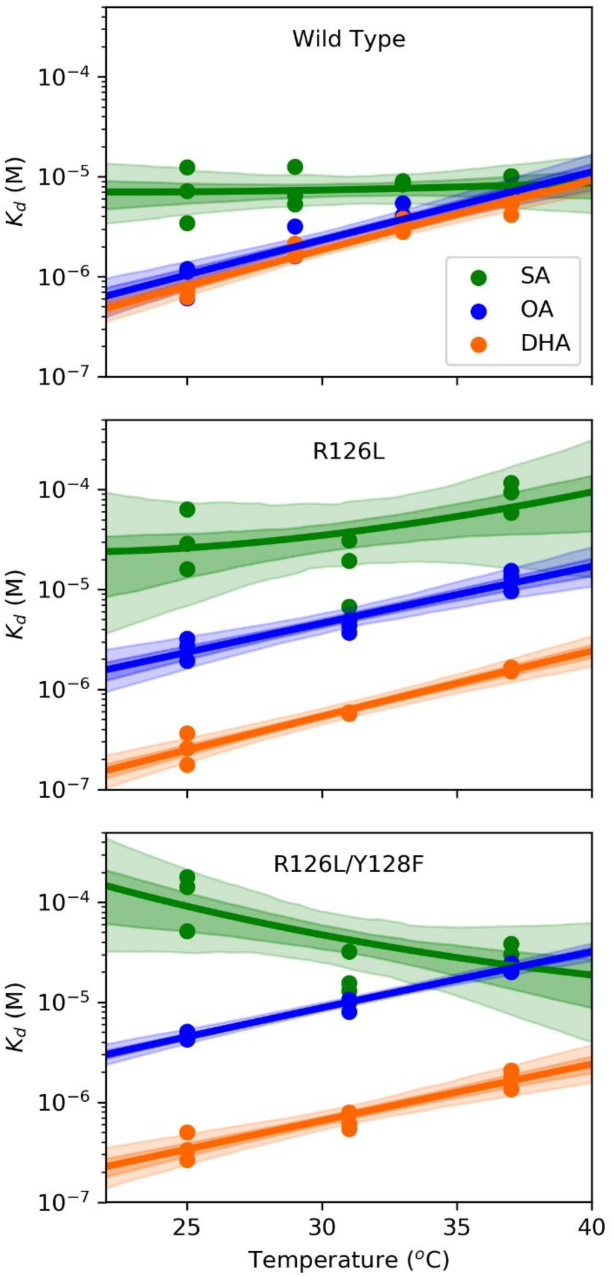


**Figure S5.** Van't Hoff Analysis of MST binding data for wt, R126L, and R16L/Y128F FABP7 show fatty acid binding is temperature dependent.

a)

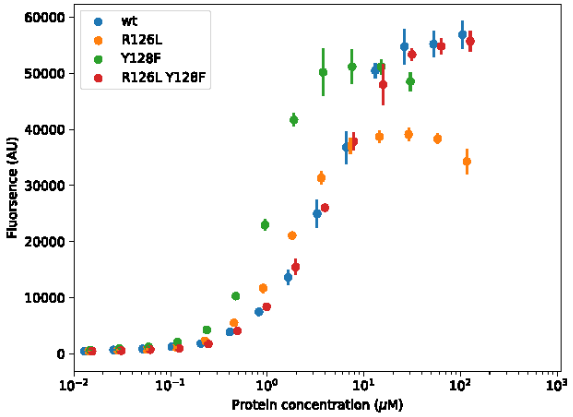


b)

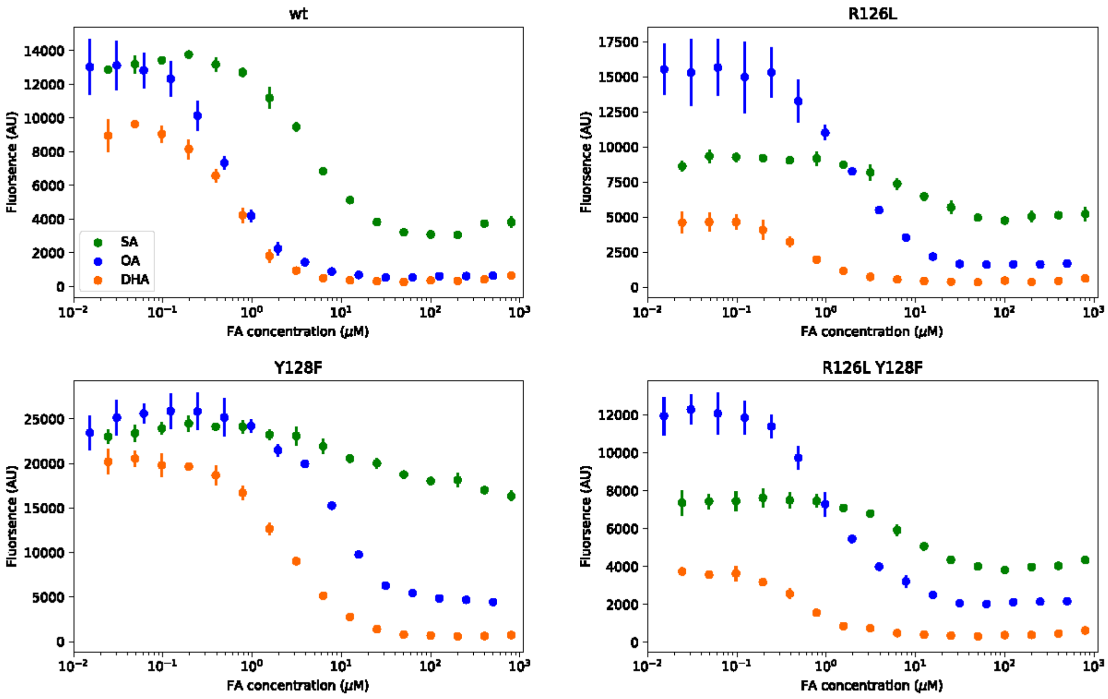


c)

| (μM) | $Kd_{ANS}$ | $EC_{50(OA)}$ | $Ki_{OA}$ | $EC_{50(SA)}$ | $Ki_{SA}$ | $EC_{50(DHA)}$ | $Ki_{DHA}$ |
|---|---|---|---|---|---|---|---|
| FABP7-wt | 4 | 0.57 | 0.253333 | 4 | 1.777778 | 0.57 | 0.253333 |
| FABP7-R126L | 1.8 | 1.8 | 0.476471 | 8 | 2.117647 | 0.488 | 0.129176 |
| FABP7-Y128F | 1 | 7.84 | 1.306667 | 20 | 3.333333 | 3.04 | 0.506667 |
| FABP7-R126L/Y128F | 4.5 | 1.06 | 0.502105 | 6 | 2.842105 | 0.556667 | 0.263684 |

**Figure S6.** a) The ANS-protein binding curves used to determine ANS dissociation constant b) ANS displacement assay curves, concentration of ANS is set to 1μM, the halfway point of this curve corresponds to the $EC_{50}$ of OA (blue), DHA (orange) and SA (green) c) shows the table of vaues measured for each curve and the associated calculated $K_i$ value from $K_i = EC_{50}/(1+[ANS]/K_{D, ANS})$

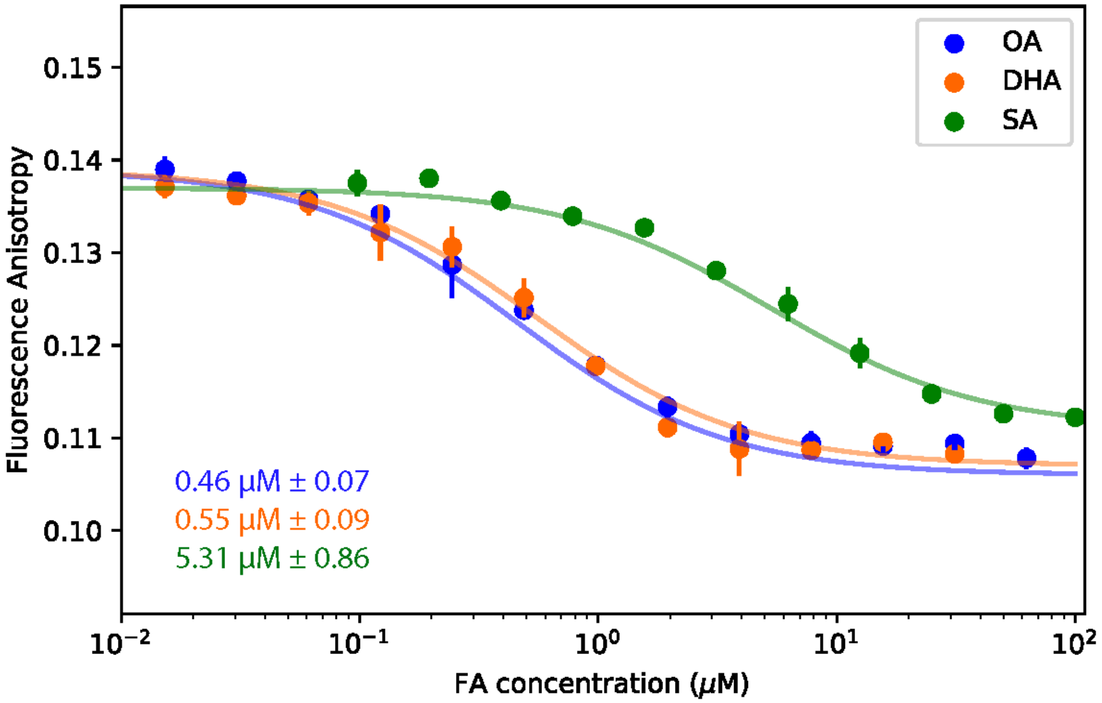


**Figure S7.** Fluorescence anisotropy for increasing concentrations of OA and DHA bound to FABP7. Upon Ligand binding we see a decrease in fluorescence anisotropy. The dissociation constants determined for OA (blue), DHA (orange), and SA (green) are shown in the bottom left of the graph.

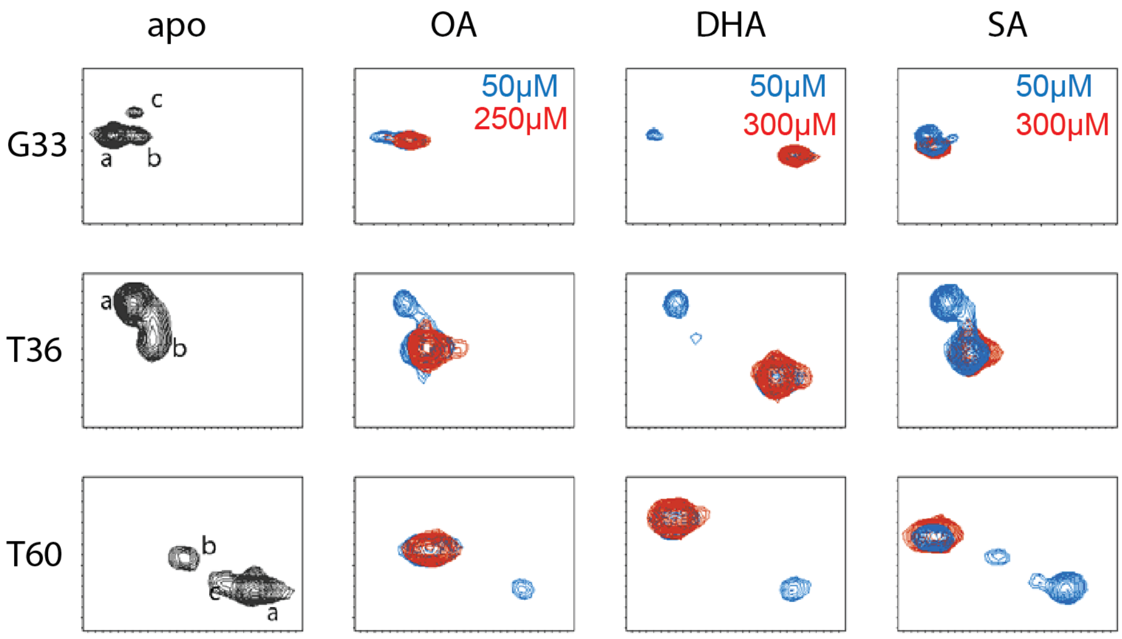


**Figure S8**. Residues which show multiple peaks in the $^{1}H$-$^{15}N$ HSQC of the apo-wtFABP7. Titration of ligand shows all three fatty acids (DHA, OA, SA) are all in slow exchange and the multiple peaks observed collapse to a single peak upon ligand saturation.

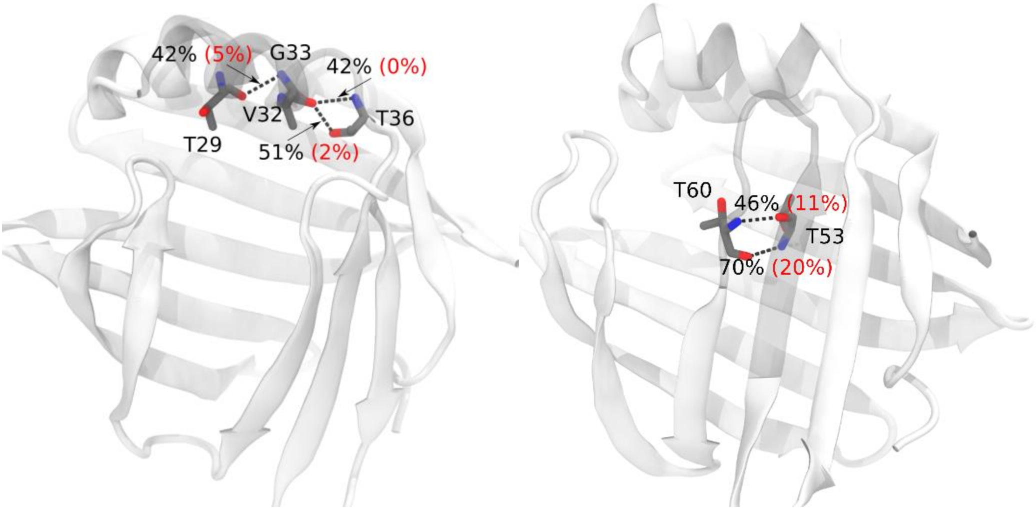


**Figure S9.** H3 and βD unfolded states are associated with reduced intra protein hydrogen bonds. Native (black) and unfolded state (red) occupancies are labelled next to dashed lines denoting hydrogen bonds.

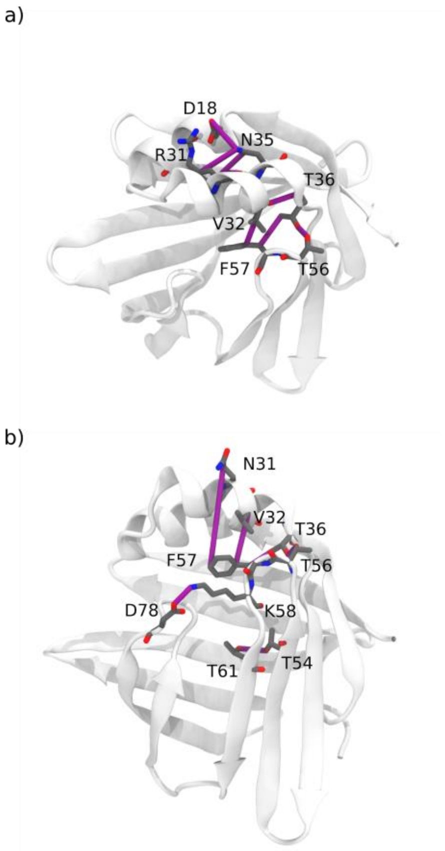


**Figure S10.** Unfolding of H3 and gap region result from loosening of contacts between the βCD turn and H3 or βEF turn. Input contact features (represented as purple cylinders) that elongate upon unfolding for the a) H3-unfolded state and b) gap-unfolded state. The unfolding of these regions is primarily driven by the presence or absence of contacts between the βCD turn and H3 or the βEF turn.

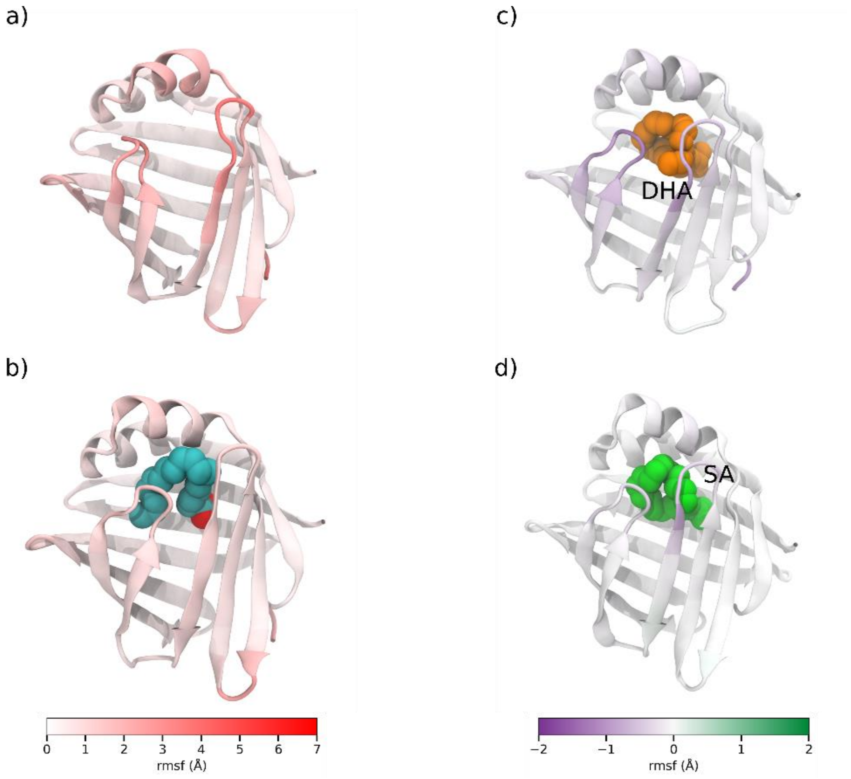


**Figure S11.** rmsf differences plots reveal that the βCD and βEF turns are more flexible for SA and DHA compared to OA with DHA having the largest difference. Absolute Cα rmsf for a) apo FABP7 and b) FABP7-OA. Cα rmsf difference between OA and c) DHA and d) SA.

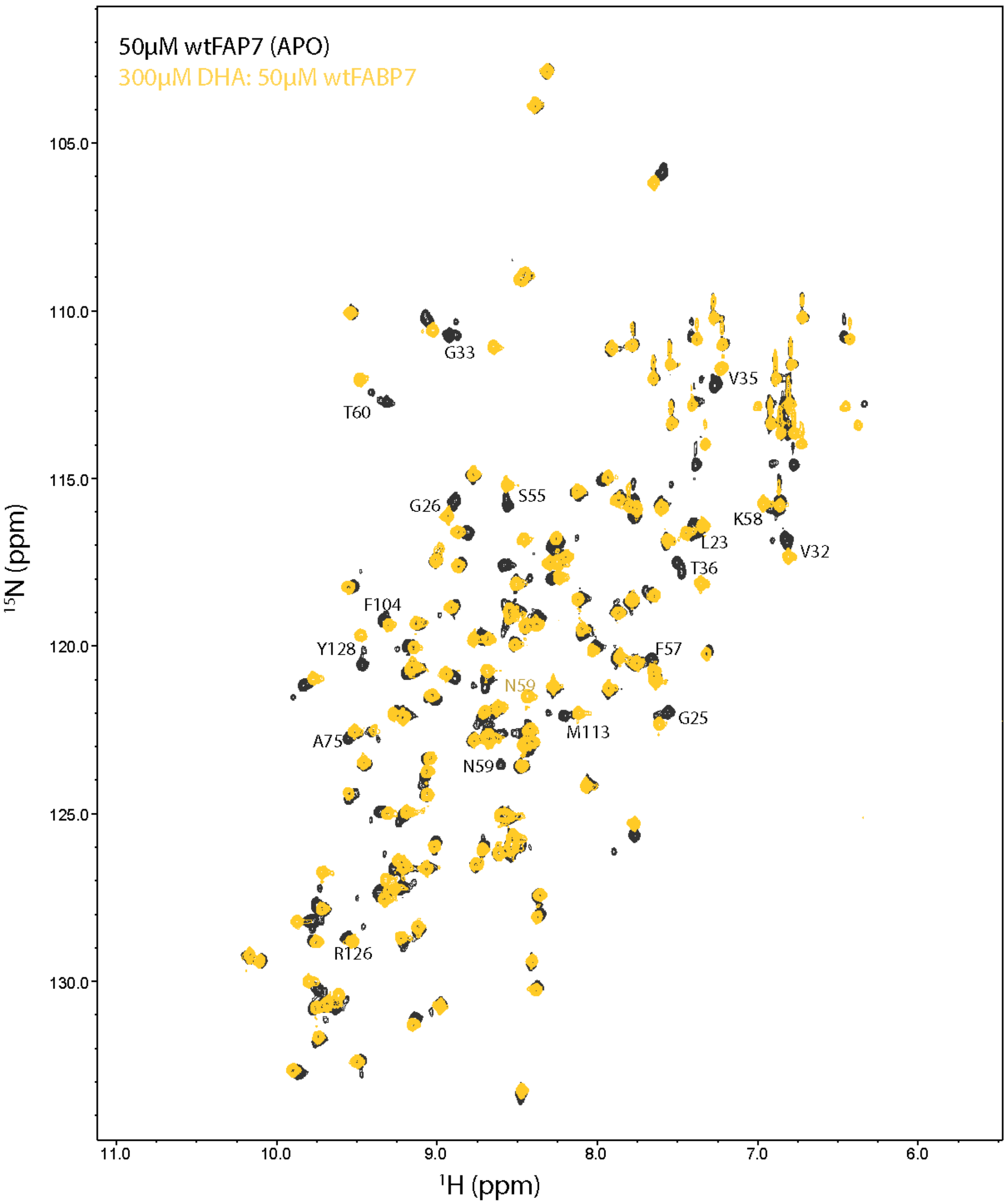


**Figure S12.** $^{1}$H-$^{15}$N HSQC showing 50µM apo-wt FABP7 (gray) and 50µM holo-wtFABP7 saturated with 300µM DHA (yellow). Peaks of importance are labeled on spectra. Assignment was imposed with literature values (13).

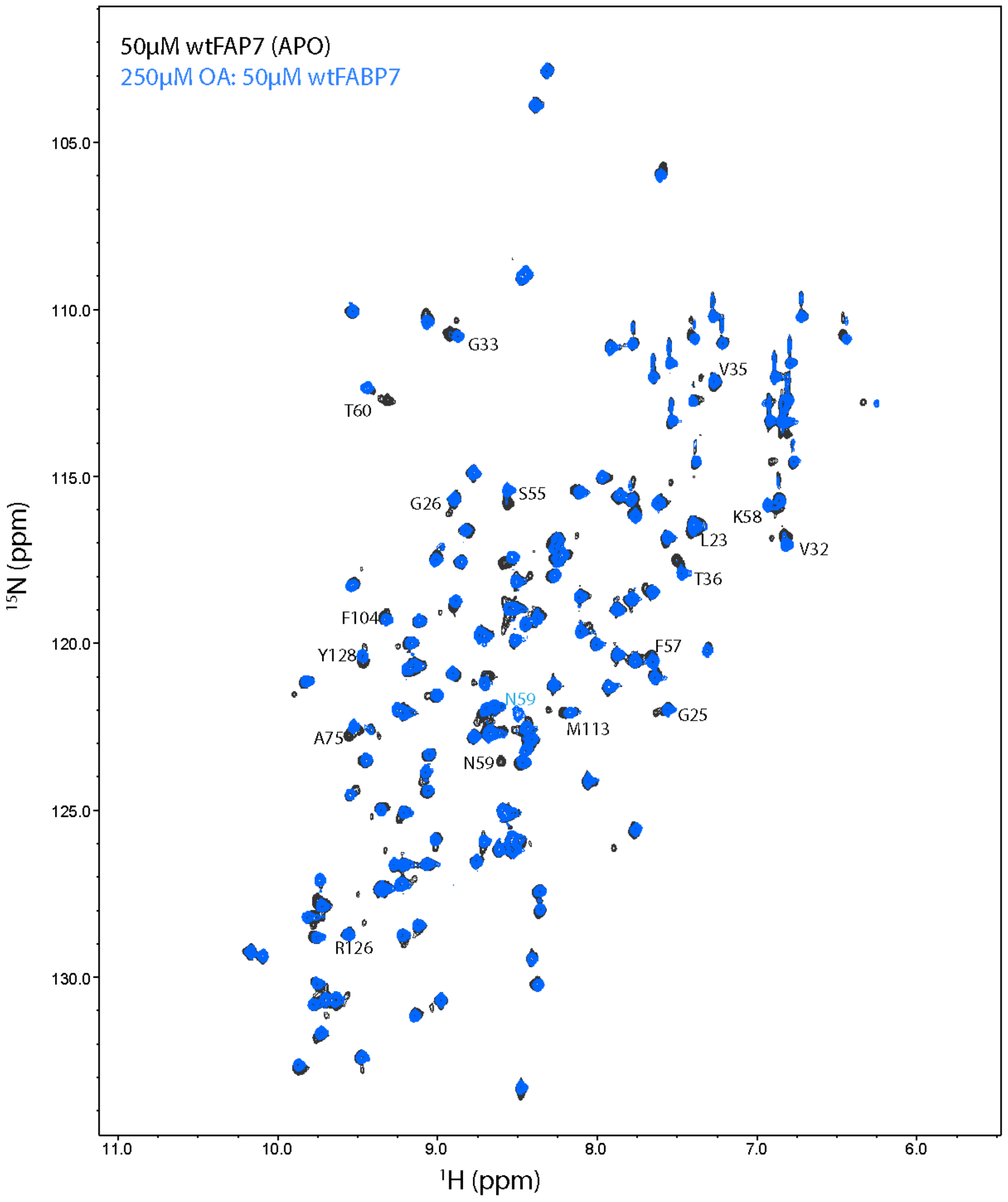


**Figure S13.** $^{1}$H-$^{15}$N HSQC showing 50μM apo-wt FABP7 (gray) and 50μM holo-wtFABP7 saturated with 250μM OA (blue). Peaks of importance are labeled on spectra. Assignment was imposed with literature values (13).

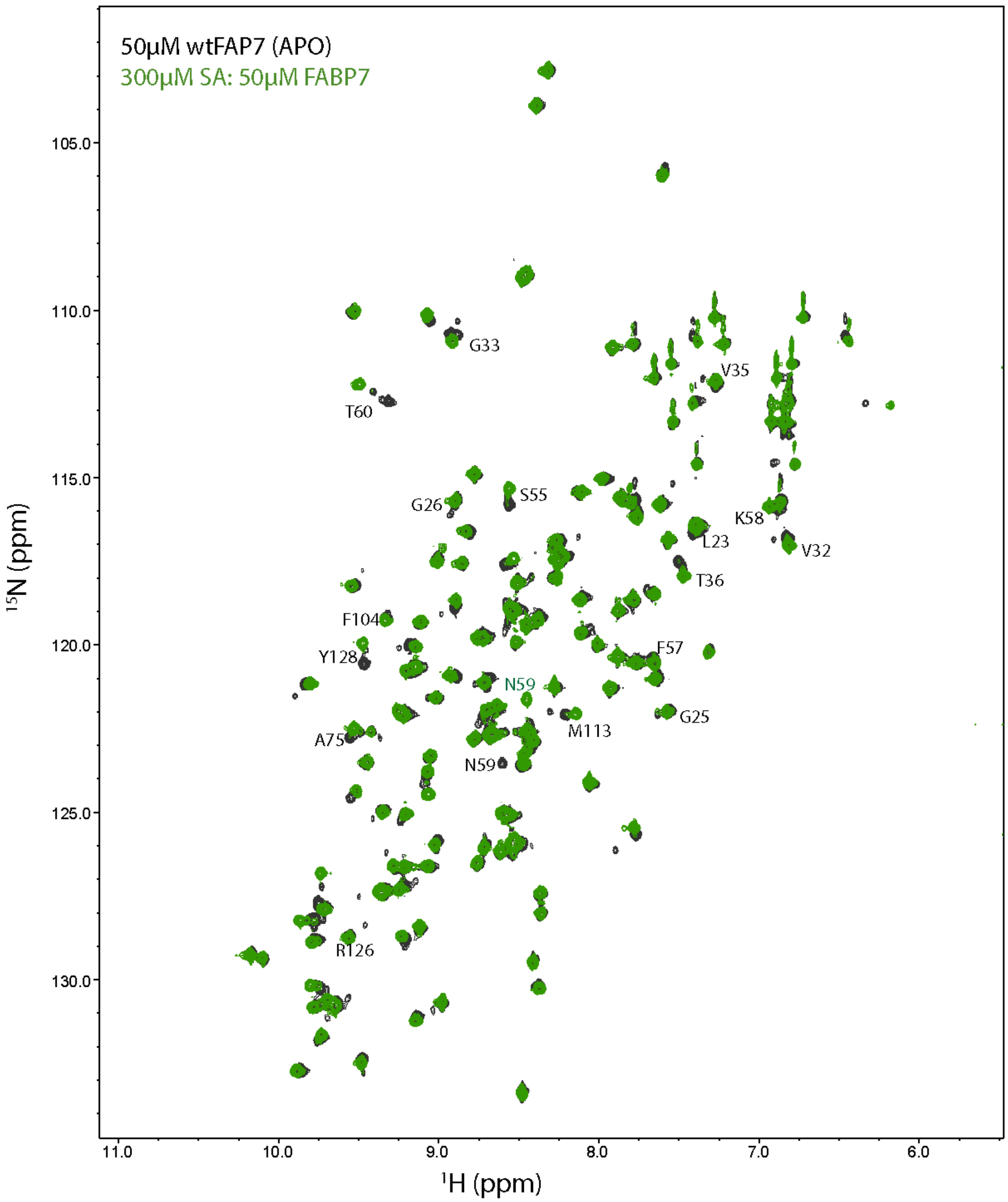


**Figure S14**. $^{1}H$-$^{15}N$ HSQC showing 50μM apo-wt FABP7 (gray) and 50μM holo-wtFABP7 saturated with 300μM SA (green). Peaks of importance are labeled on spectra. Assignment was completed by peak following and comparison to literature values (13).

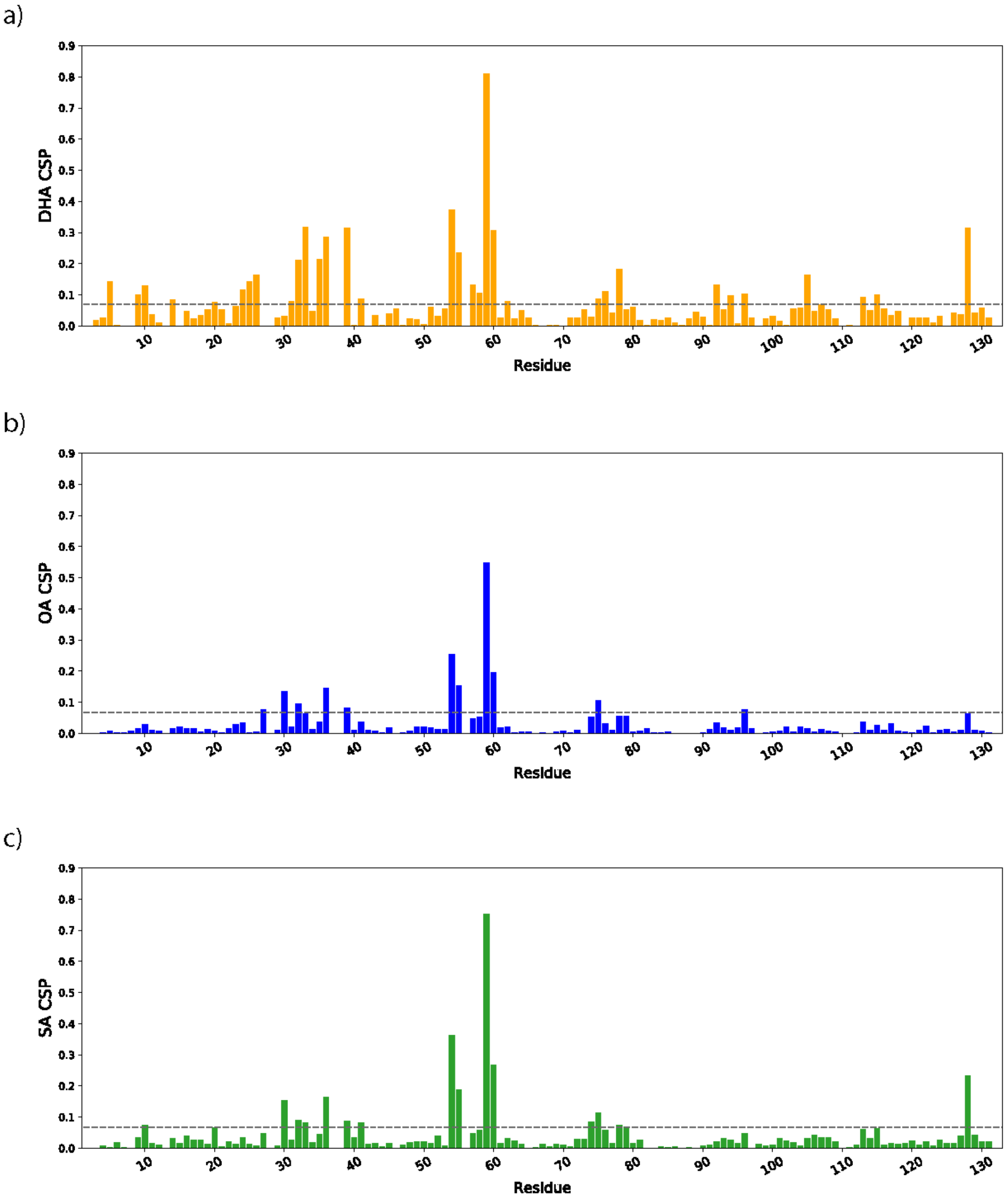


**Figure S15.** CSP for each residue associated with fatty acid binding of a) DHA, b) OA, and c) SA.

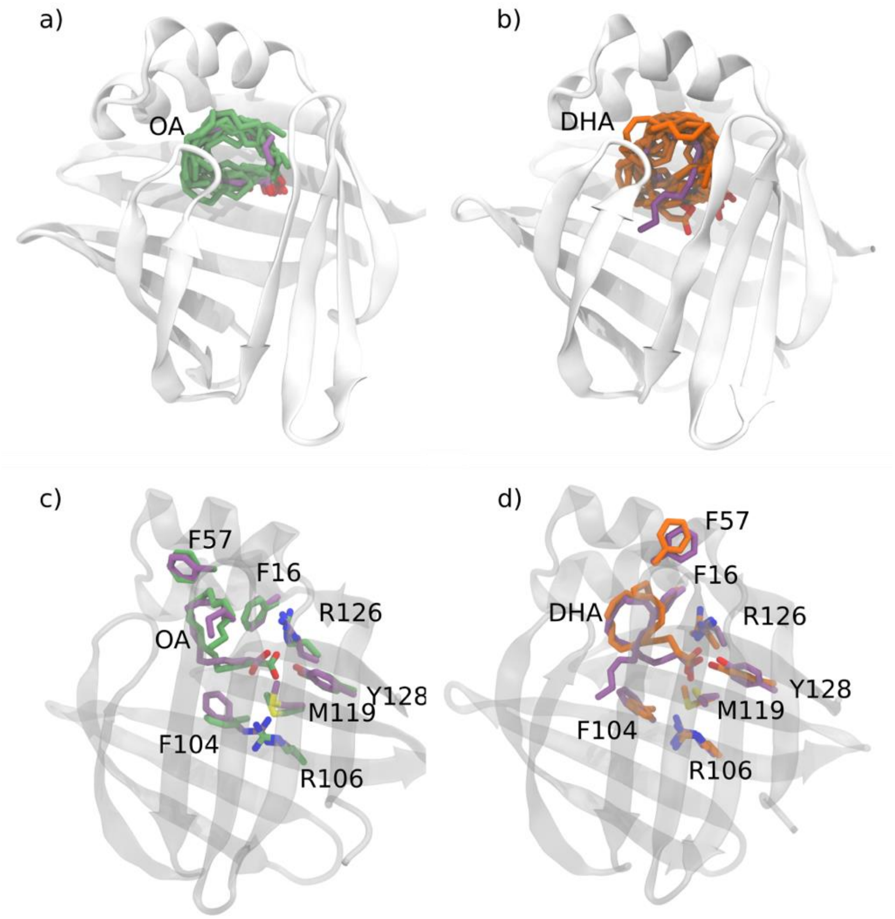


**Figure S16.** Superposition of HMM-sampled structures from most populous state for a,c) OA and b,d) DHA onto crystal structures of FABP7 bound to OA (PDB:1fe3) or DHA (PDB:1fdq). For a) and b) 10 structures were randomly sampled to provide a qualitative estimate of the ligand binding pose heterogeneity, while c) and d) show the binding pocket for a single HMM-sampled structure.

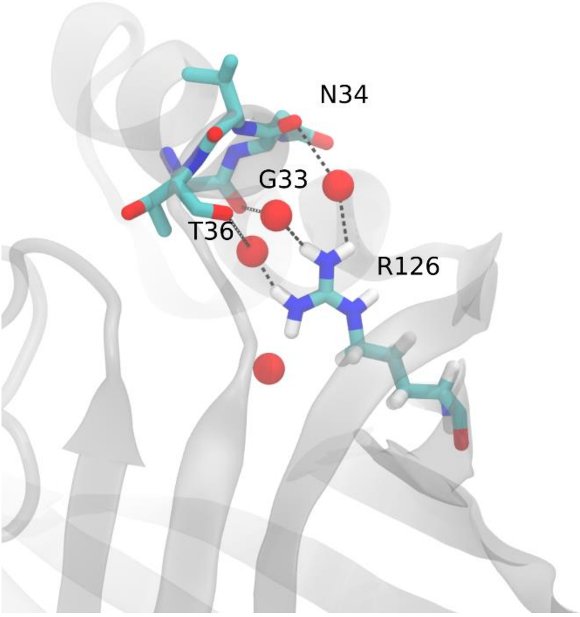


**Figure S17.** Water-mediated hydrogen bonds (water oxygen atoms are represented as red spheres) between R126 and G33, N34, and T36 are lost upon mutation of R126 leading to greater H3 instability.

binding protein (FABP7) in its apo form and the holo forms binding to DHA, oleic acid, linoleic acid and elaidic acid. *Biomol NMR Assign*. 3:89–93.
